# ER cholesteryl ester phase separation underlies switch-like cholesterol sensing

**DOI:** 10.64898/2026.08.01.740796

**Authors:** Mehdi Zouiouich, Apoorva Bhapkar, Jennica Träger, Murphy McDermott, Sonny Yde, Sami Kchir, Fabienne Foufelle, Mohyeddine Omrane, Abdou Rachid Thiam

**Affiliations:** Laboratoire de Physique de l’École Normale Supérieure, ENS, Université PSL, CNRS Sorbonne Université, Université Paris-Diderot, Sorbonne Paris Cité, Paris, France; Sorbonne Université, INSERM, UMRS1166, Paris, France

## Abstract

Cellular lipid homeostasis requires mechanisms that detect subtle changes in lipid abundance and trigger rapid, coordinated responses. The INSIG/SCAP/SREBP2 pathway provides a central feedback system linking endoplasmic reticulum (ER) cholesterol levels to the transcriptional control of cholesterol genes, yet the origin of its remarkable cooperative switch mechanism remains unclear. Here we identify cholesterol esterification as a physical mechanism that amplifies sterol sensing and generates switch-like pathway regulation. We show that cholesteryl oleate (CE), produced by SOAT1 and opposed by NCEH1-mediated hydrolysis, undergoes a cooperative phase transition within the ER membrane to form transient CE-rich domains. These lipid assemblies create a threshold-dependent platform that concentrates SCAP and promotes formation of the SCAP-INSIG retention complex, thereby coupling ER lipid organization to SREBP inhibition. Because CE domain formation is nucleation-driven, variations in cholesterol availability are converted into an abrupt transition between distinct membrane states. Perturbing CE metabolism uncouples cholesterol abundance from pathway activity: SOAT1 inhibition prevents CE domain formation, releases SCAP from the ER, and activates SREBP2 despite cholesterol sufficiency, whereas NCEH1 inhibition stabilizes the domains and reduces SREBP2 activation under cholesterol-limiting conditions. Thus, the balance between esterification and hydrolysis determines a membrane physical state that serves as the functional output sensed by the cholesterol regulatory machinery. Our findings reveal ER lipid phase transitions as a general principle for creating ultrasensitive control in cellular homeostasis and establish cholesterol esterification as an active regulatory process rather than a passive storage pathway.

## INTRODUCTION

Cholesterol homeostasis relies on a feedback circuit centered on the endoplasmic reticulum (ER), where the INSIG/SCAP-SREBP pathway couples membrane sterol content to transcriptional control of cholesterol synthesis and uptake^1,2^. When ER cholesterol falls below ∼5 mol%^3–5^, SCAP escorts SREBP to the Golgi for proteolytic activation, inducing the expression of genes required for cholesterol biosynthesis and LDL uptake^6,7^. As ER cholesterol rises, cholesterol binding to the sterol-sensing domain of SCAP induces conformational changes that promote its association with INSIG proteins, retaining the SCAP-SREBP complex in the ER and shutting down SREBP processing^3,8,9^. Oxysterols reinforce this feedback by binding INSIG^8,10^ and simultaneously potently activating sterol O-acyltransferase 1 (SOAT1), thereby coupling cholesterol sensing to cholesterol esterification^11,12^ (Fig. 1A). Recent cryo-electron microscopy studies have revealed the structural basis of cholesterol recognition by SCAP and its interaction with INSIG^10,13^, establishing a molecular framework for this regulatory switch.

**Figure 1.**
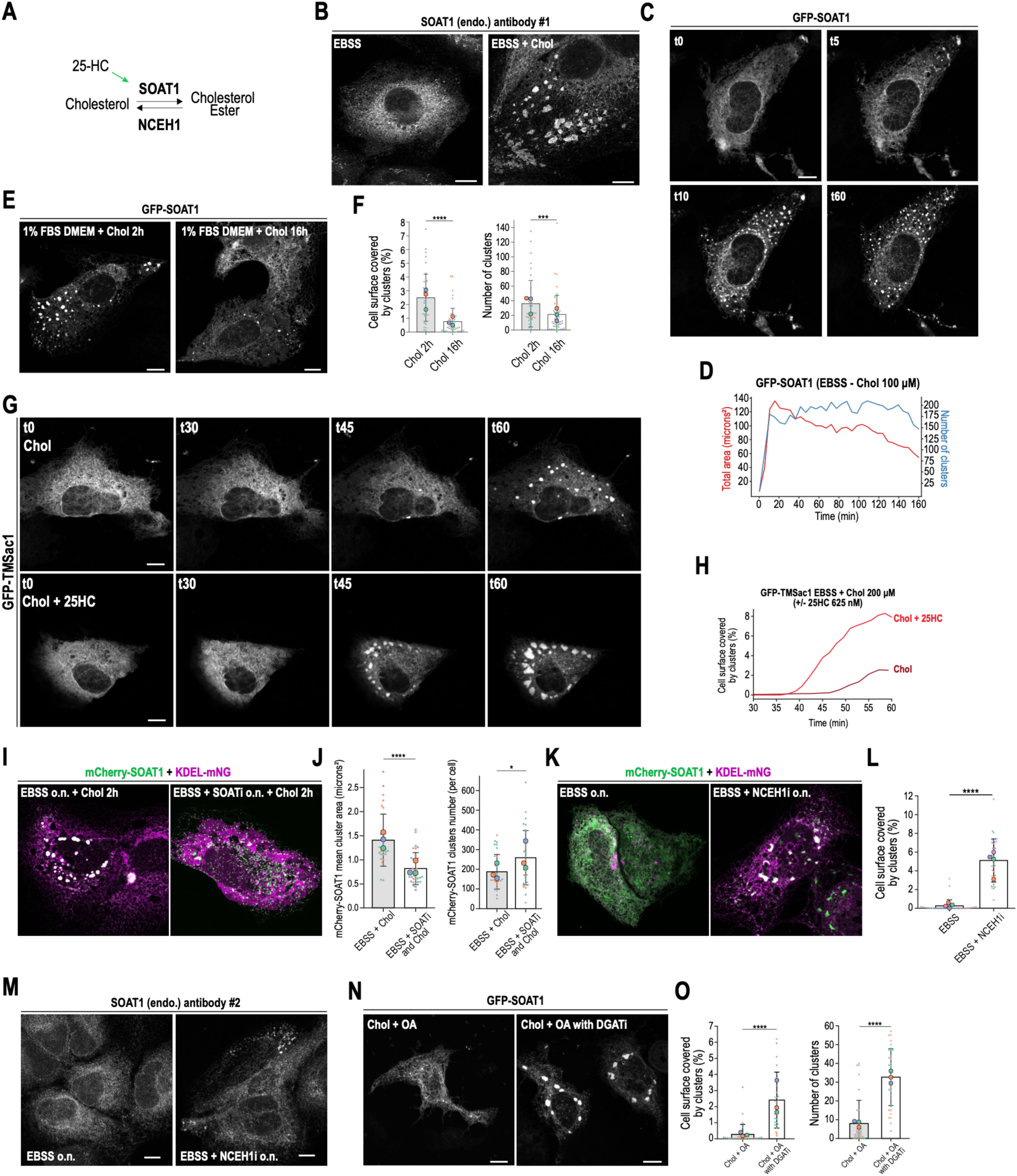
Cholesteryl esters drive the formation of transient ER microdomains. **(A)** Scheme showing the role of SOAT1 and NCEH1 related to cholesteryl esters. 25-hydroxycholesterol (25-HC) stimulates SOAT1 activity. **(B)** Images of immunofluorescence (IF) experiments performed in Huh7 cells against SOAT1 (antibody #1) in starved cells (left, EBSS) and cholesterol-loaded (100 µM for 2 hours) cells representing an extreme condition (right, EBSS + Chol). **(C)** Time-lapse images of Huh7 cells overexpressing GFP-SOAT1. Cells were in EBSS and fed with 100 µM of cholesterol (t0 corresponds to the time of cholesterol addition). **(D)** Quantification of the total area (in red) and the number (in blue) of microdomains from the time-lapse shown in (C). **(E)** Images of Huh7 cells overexpressing GFP-SOAT1 grown in DMEM + 1% FBS. Cholesterol was added at 100 µM for either 2 hours (left) or 16 hours (right). **(F)** Quantification of (E) showing the percentage of cell surface covered by clusters (left) and the number of clusters (right). Data collected from 3 biological replicates (n=42 Chol 2 hours; n=49 Chol 16 hours). A Mann-Whitney U test was performed for statistical analysis. **(G)** Timelapse images of Huh7 cells overexpressing GFP-TM Sac1. Cells were starved overnight in EBSS, then fed with 200 µM of cholesterol (upper panels) or 200 µM of cholesterol + 625 nM of 25-HC (bottom panels). **(H)** Quantification of the percentage of cell surface covered by microdomains from time-lapses shown in (G). **(I)** Merged images of Huh7 cells overexpressing mCherry-SOAT1 (in green) and KDEL-mNeonGreen (mNG) (in magenta). Cells were starved overnight in EBSS only (left) or in EBSS with SOAT inhibitor (SOATi) (right), then fed with cholesterol at 100 µM for 2 hours. **(J)** Quantification of (I) showing the mean size (left) and number of clusters (right). Data collected from 3 biological replicates (n=31 EBSS + Chol 2h; n=32 EBSS + SOATi + Chol 2 hours). A Mann-Whitney U test was performed for statistical analysis. **(K)** Merged images of Huh7 cells overexpressing mCherry-SOAT1 (in green) and KDEL-mNeonGreen (in magenta). Cells were starved overnight (o.n.) in EBSS only (left) or in the presence of NCEH1 inhibitor (NCEH1i) (right). **(L)** Quantification of (K) showing the percentage of the cell surface covered by microdomains. Data collected from 4 biological replicates (n=33 EBSS; n=32 EBSS + NCEH1i). A Mann-Whitney U test was performed for statistical analysis. **(M)** Images of IF studies performed in Huh7 cells against SOAT1 starved in EBSS alone (left) or in the presence of NCEH1 inhibitor (right). **(N)** Images of Huh7 cells overexpressing GFP-SOAT1 starved in EBSS overnight alone (left) or in the presence of DGAT inhibitors (DGATi) (right), then fed with cholesterol (50 µM) + oleic acid (OA) (50 µM) for 2 hours. **(O)** Quantification of (N) showing the percentage of cell surface covered by microdomains (left) and their number (right). Data collected from 3 biological replicates (n=34 Chol + OA; n=36 Chol + OA with DGATi). A Mann-Whitney U test was performed for statistical analysis. Scale bars represent 10 µm.

Despite these advances, fundamental properties of the INSIG/SCAP-SREBP system remain unresolved^3,5^. How is cholesterol sensed with such precision? Although ER cholesterol is maintained near ∼5 mol%, synthesis, uptake, and inter-organelle transport continuously perturb its levels^14–16^. What defines this specific critical concentration is unknown. If SCAP would respond to the average cholesterol concentration of the ER membrane, fluctuations would generate persistent noise and inappropriate activation of the pathway. Instead, SREBP regulation is remarkably robust^3^, suggesting that SCAP may not directly sample bulk ER cholesterol. Second, how is an ultrasensitive response generated? SREBP processing transitions from fully active to fully repressed over this exceptionally narrow cholesterol concentration range, corresponding to an apparent Hill coefficient of approximately 3-7^3^. Such switch-like behavior cannot be explained by a simple cholesterol equimolar binding to SCAP, while the known oligomeric organization of SCAP provides only a partial explanation^2^. The origin of this emergent cooperativity remains unknown. Lastly, what determines the kinetics of the switch? Cholesterol loading immobilizes SCAP in the ER within minutes, with retention essentially complete after tens of minutes^8,17,18^. By contrast, although cholesterol depletion rapidly initiates SCAP release, complete restoration of SREBP signaling requires several hours because SCAP mobilization is followed by sequential proteolytic processing, nuclear translocation, and transcriptional activation^2,8,18–20^. The mechanisms that establish rapid retention yet delay complete pathway reactivation remain unclear.

Together, these observations point to a model in which SCAP regulation depends not on bulk ER cholesterol concentration but on a localized membrane environment that filters transient fluctuations, amplifying cooperativity beyond what molecular interactions alone can produce.

Cholesterol ester (CE) is a cholesterol product/substrate produced in the ER by SOAT1 and hydrolyzed by the Neutral Cholesterol Ester Hydrolase (NCEH1)^21^ (Fig.1A), AADACL1, belonging to the arylacetamide deacetylase (AADAC) protein family, which is a small group of ER-associated serine hydrolases. CE is widely thought to serve merely as a cholesterol storage molecule in lipid droplets (LDs), where it adopts highly ordered crystalline phases capable of altering membrane protein distribution ^22–25^. We examined whether CE is involved in SREBP regulation.

## RESULTS

### Cholesterol esterification drives SOAT1 clustering into transient ER microdomains

To investigate the spatial organization of cholesterol-handling machinery in the ER, we monitored endogenous SOAT1 localization by immunofluorescence in Huh7 hepatocellular carcinoma cells. To establish a cholesterol-depleted state in which ER cholesteryl ester (CE) stores are minimized and SOAT1 activity is reduced, cells were maintained overnight in low serum (1% FBS) or serum-free Earle’s balanced salt solution (EBSS). Under these conditions, SOAT1 displayed a diffuse ER distribution (Fig. 1B). In contrast, acute cholesterol supplementation (100 µM cholesterol delivered by methyl-β-cyclodextrin, MβCD) induced the rapid redistribution of SOAT1 into discrete, highly concentrated ER structures, which we termed microdomains (Fig. 1B and Fig. S1A). Similar microdomain formation was observed in HeLa cells, indicating that this response is not restricted to hepatocytes (Fig. S1B).

To define the dynamics of microdomain formation, we performed live-cell imaging of Huh7 cells expressing GFP-SOAT1. Following cholesterol addition (100 µM), microdomains nucleated rapidly within 5–10 minutes, followed by a plateau phase characterized by continuous fusion and fission events (Fig. 1C-D, Fig. S1C, Video S1), consistent with a cooperative nucleation and growth assembly process. Lower cholesterol supplementation (50 µM) also induced SOAT1 clustering, although with reduced domain size, demonstrating a dose-dependent response to cholesterol availability (Fig. S1D-E). Remarkably, microdomains were transient and largely disappeared by 16 hours after cholesterol loading (Fig. 1E-F, Video S2), suggesting that they represent dynamic transient ER states rather than stable assemblies.

The kinetics of microdomain formation were strongly influenced by the metabolic state of the cell. Cholesterol loading in cells maintained in 1% FBS resulted in more rapid and extensive microdomain formation, followed by faster dissolution compared with EBSS-incubated cells (Fig. S1F-G, Video S3). Furthermore, a cholesterol pulse-wash experiment (2 h cholesterol loading followed by transfer to DMEM containing 10% FBS) halted microdomain expansion and promoted their dissolution (Fig. S1H-I, Video S4), indicating that continued cholesterol delivery is required to sustain these structures. During dissolution, lipid droplets (LDs) formed and expanded concomitantly with microdomain disappearance (Fig. S1H-I), suggesting that cholesterol redistribution into LDs may contribute to microdomain dissolution.

Microdomains were largely negative for the autophagic marker LC3B, including during pulse-wash conditions that accelerated their disappearance (Fig. S1J-K, Video S5), distinguishing them from LC3B-positive reticulophagy intermediates and ER stress bodies^26^.

Because SOAT1 catalyzes cholesterol esterification, we next tested whether increased SOAT1 activity was sufficient to accelerate microdomain formation. In cells without SOAT1 overexpression, cholesterol addition to EBSS-starved cells induced SOAT1 clustering only after approximately one hour, using the ER transmembrane marker Sac1, in contrast to 5-10min in SOAT1 overexpression (Fig.1C). This suggests that SOAT1 activity controls the kinetics of microdomain formation. Consistently, co-addition of 25-HC, which enhances endogenous SOAT1 activation, increased microdomain formation in response to cholesterol loading (Fig. 1G-H, Video S6-S7). A delayed response was similarly observed when cells were maintained in 1% FBS rather than EBSS (Fig. S1L-M, Video S8), further demonstrating that cellular lipid state regulates microdomain dynamics.

Microdomains colocalized with the ER lumen marker KDEL-mNeonGreen (Fig. S1N), confirming their ER localization. Importantly, pharmacological inhibition of SOAT1 before cholesterol supplementation prevented microdomain formation (Fig. 1I-J), demonstrating that SOAT1 catalytic activity is required. We therefore asked whether increasing ER CE levels through inhibition of CE hydrolysis would be sufficient to induce these structures. Inhibition of the CE hydrolase NCEH1 for 1 hour prior to nutrient deprivation and throughout the deprivation period resulted in robust microdomain formation without exogenous cholesterol addition (Fig. 1K-L). This effect was observed with endogenous SOAT1 (Fig. 1M) and was further enhanced by cholesterol supplementation (Fig. S2A-C).

Together, these results demonstrate that ER microdomains are driven by net CE accumulation and that their formation is controlled by the balance between SOAT1-mediated cholesterol esterification and NCEH1-mediated CE hydrolysis. Thus, SOAT1 and NCEH1 establish a CE metabolic circuit that regulates the emergence of cholesterol-responsive ER membrane domains.

### Microdomain formation and lifetime are regulated by DGATs

Microdomain disappearance coincided with LD formation and growth (Fig. S1H-I), suggesting that LD biogenesis may provide a mechanism for resolving ER CE accumulation. We previously showed that CE incorporation into LDs requires triacylglycerol (TAG) synthesis by diacylglycerol O-acyltransferase (DGAT) enzymes, and that the TAG/CE ratio governs LD growth and internal organization^23,24^. We therefore asked whether TAG availability influences ER microdomain dynamics.

Co-supplementation of cholesterol with oleic acid (OA), which stimulates TAG synthesis and LD expansion, strongly suppressed microdomain formation (Fig. 1N-O). Similarly, overexpression of either DGAT1 or DGAT2 markedly reduced the appearance of microdomains (Fig. S2D-E). Conversely, pharmacological inhibition of DGAT activity prior to cholesterol loading enhanced microdomain formation (Fig. 1N-O), consistent with impaired CE transfer into LDs^23^ and increased retention of CE within the ER membrane.

Together, these results reveal a three-way balance controlling ER CE organization: SOAT1 promotes microdomain formation by generating CE, NCEH1 counteracts accumulation through CE hydrolysis, and DGAT enzymes limit microdomain persistence by promoting CE sequestration into LDs. Thus, ER microdomain abundance and lifetime are determined by the integrated activity of CE synthesis, hydrolysis, and lipid storage pathways.

This model provides a mechanistic explanation for the transient behavior of microdomains observed by live imaging. Following cholesterol loading, initial SOAT1-dependent CE production promotes rapid domain nucleation, whereas subsequent DGAT-dependent LD expansion progressively redistributes CE away from the ER membrane, leading to domain dissolution. The kinetics of microdomain disappearance therefore mirror the timescale of LD biogenesis following cholesterol excess^23^.

### Cholesteryl oleate drives lipid phase separation in ER-mimetic and native membranes

We next asked whether CE accumulation within the ER membrane bilayer is sufficient to drive lipid phase separation. To modulate ER CE content across a broad physiological range, we established four complementary conditions: (1) overnight EBSS to deplete CE; (2) EBSS followed by a 2-hour MβCD-cholesterol pulse to stimulate CE synthesis; (3) NCEH1 inhibition during nutrient deprivation to promote CE accumulation by blocking hydrolysis; and (4) SOAT1 inhibition during cholesterol loading to prevent CE synthesis despite cholesterol availability. Together, these perturbations provided a controlled framework to define the relationship between ER CE content and membrane organization. ER fractions were isolated from each condition by density-gradient ultracentrifugation.

ER fractions enriched in calnexin and lacking LD markers (Fig. S2F) were subjected to lipidomic profiling. Nutrient deprivatino selectively reduced cholesteryl oleate (CE 18:1), whereas polyunsaturated CE species were largely preserved (Fig. 2A). Conversely, cholesterol supplementation strongly increased CE 18:1, consistent with the substrate preference of SOAT1 for oleoyl-CoA^27,28^. SOAT1 inhibition markedly reduced ER CE levels, with the strongest effect on CE 18:1, whereas NCEH1 inhibition promoted accumulation of CE 18:1 and additional saturated CE species (Fig. 2A). These data demonstrate that cholesteryl oleate is the major CE species dynamically regulated within the ER membrane through opposing SOAT1-dependent synthesis and NCEH1-dependent hydrolysis.

**Figure 2.**
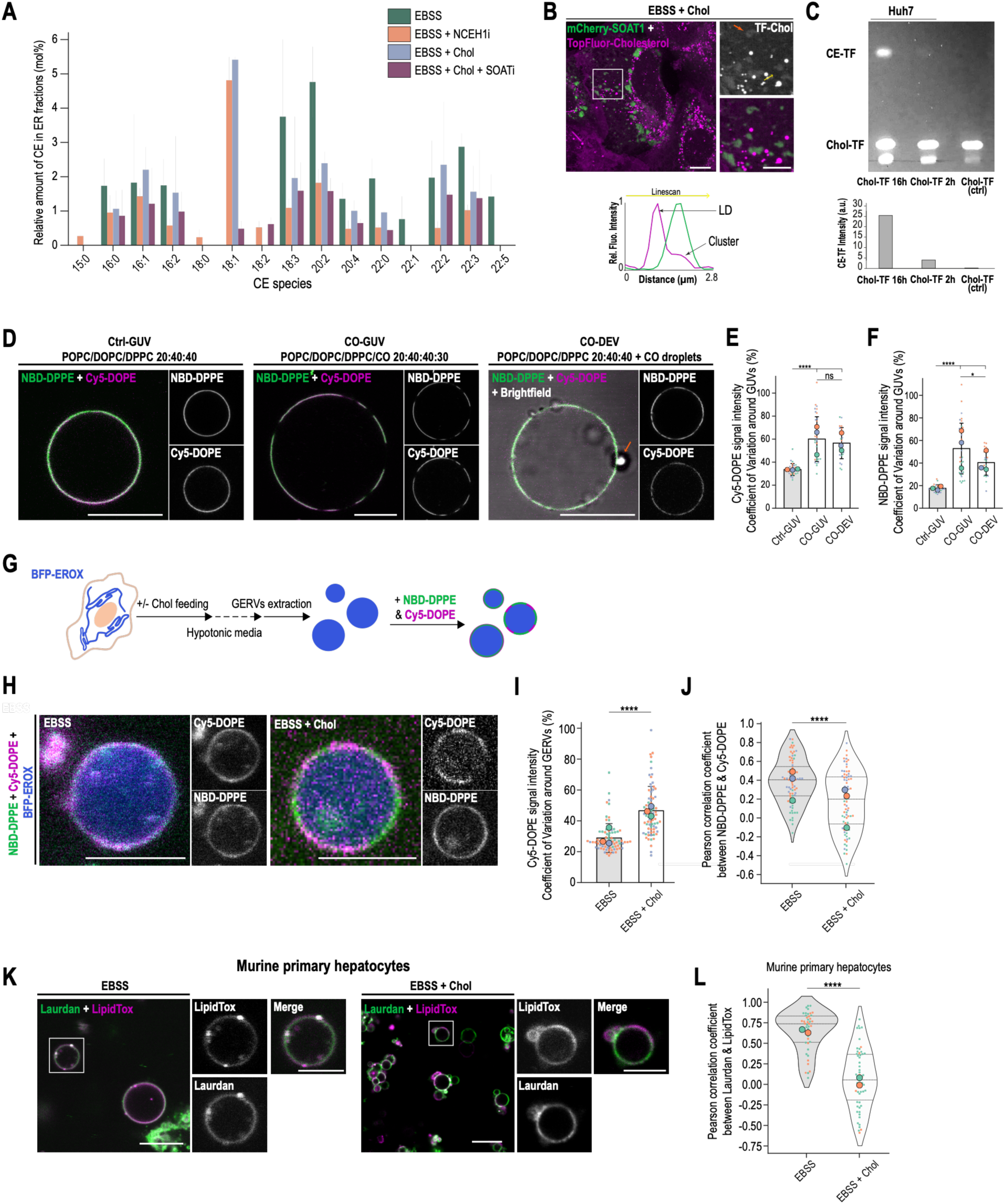
Cholesteryl esters within the ER membrane induce lipid phase separation. **(A)** Quantification of the relative amount of cholesteryl esters (CE) within the ER fraction after density-based organelle purification. Huh7 cells were starved overnight with the NCEH1 inhibitor (in orange), the SOAT inhibitor (in purple), or no drug (in green). Cholesterol was added at 200 μM for 2 hours under two conditions: with the SOAT inhibitor (purple) or without the inhibitor (blue). Data collected from 3 biological replicates. **(B)** Merge image of Huh7 cells overexpressing mCherry-SOAT1 (in green) fed with a mix of cholesterol (150 μM) + TopFluor-Cholesterol (50 μM) (in magenta) for 2 hours. The orange arrow indicates the TF-Chol signal within the microdomain. The yellow arrow represents the linescan shown on the right panel. **(C)** Thin layer chromatography image revealed by fluorescence. Huh7 cells were incubated with cholesterol and TopFluor-cholesterol (Chol-TF) either overnight (16 hours) or for 2 hours. Afterward, cell extracts were collected and analyzed. The control sample, with TopFluor-cholesterol deposited, is shown in the left column. **(D)** Merged images of GUVs labeled with NBD-DPPE (in green) and Cy5-DOPE (in magenta). GUVs are made with POPC/DOPC/DPPC (20:40:40) (Ctrl-GUV, left panel) or + 30 mol% of cholesteryl oleate (CO) (CO-GUV, middle panel). CO-DEV corresponds to Ctrl-GUV with added artificial CO droplets (right panel). **(E-F)** Quantification of the coefficient of variation of Cy5-DOPE (E) or NBD-DPPE (F) from conditions shown in (D). Data collected from 3 independent replicates (n=34 Ctrl-GUV; n=30 CO-GUV; n=24 CO-DEV). Significant differences between groups were determined using a Kruskal-Wallis test followed by post hoc pairwise Mann-Whitney U tests with Bonferroni correction. **(G)** Representation of the main steps performed to extract and label GERVs. **(H)** Merged images of GERVs from Huh7 cells overexpressing BFP-EROX (in blue) and labeled with NBD-DPPE (in green) and Cy5-DOPE (in magenta). Cells were either starved overnight without treatment (EBSS, left) or fed with cholesterol at 200 μM for 2 hours (EBSS + Chol, right). **(I-J)** Quantification of the coefficient of variation of Cy5-DOPE signal (H) or the Pearson correlation coefficient between the NBD and Cy5 signals (J) from conditions shown in (H). Data collected from 3 biological replicates (n=79 EBSS; n=79 EBSS + Chol). A Mann-Whitney U test (I) and a Welch’s t-test (J) were performed for statistical analysis. **(K)** Merged images of GERVs from murine primary hepatocytes labeled with Laurdan (in green) and LipidTOX (in magenta), starved overnight in EBSS alone (left) or with 200 μM cholesterol for 2 hours (right). **(L)** Quantification of the Pearson correlation coefficient between the Laurdan and LipidTOX signals from conditions shown in (K). Data collected from 2 mice (n=43 EBSS; n=58 EBSS + Chol). A Mann-Whitney U test was performed for statistical analysis. Scale bars represent 10 µm; insets from (B) and (K), 5 µm.

To directly visualize CE enrichment within ER microdomains, cells were loaded with TopFluor-cholesterol, which is converted into fluorescent CE (Fig. 2C). The fluorescent signal accumulated both in LDs and in SOAT1-positive ER microdomains (Fig. 2B), demonstrating that ER CE-rich domains and LDs represent distinct but interconnected CE-containing structures.

We next tested whether cholesteryl oleate itself can induce membrane phase separation. Giant unilamellar vesicles (GUVs) reproducing the major ER lipid composition (POPC/DOPC/DPPC, 1:2:2) were supplemented with 30 mol% cholesteryl oleate. CE incorporation induced robust lateral phase separation at 37°C (Fig. 2D-F). To determine whether CE stored in LDs can redistribute into adjacent membranes, we generated droplet-embedded vesicles (DEVs) by introducing cholesteryl oleate-containing artificial LDs (aLDs) into initially homogeneous GUVs^29^. Transfer of CE from aLDs into the surrounding bilayer induced phase-separated domains indistinguishable from those generated by direct CE incorporation (Fig. 2D–F), indicating that LD-derived CE can partition into ER-like membranes and nucleate membrane organization.

To establish whether CE-dependent phase separation occurs in native ER membranes, we used giant ER vesicles (GERVs), which preserve the native lipid and protein composition of ER membranes^30^. Membrane organization was assessed using the liquid-disordered marker NBD-DPPE and the liquid-ordered marker Cy5-DOPE (Fig. 2G). GERVs derived from EBSS-treated cells displayed homogeneous probe distribution, whereas GERVs from cholesterol-loaded cells exhibited pronounced lateral segregation of Cy5-DOPE (Fig. 2H). This reorganization was quantified by an increased coefficient of variation (Fig. 2I) and reduced probe colocalization (Fig. 2J), demonstrating CE-dependent phase separation within native ER membranes. Importantly, GERVs isolated from primary mouse hepatocytes following cholesterol feeding displayed comparable membrane segregation (Fig. 2K-L), confirming the physiological relevance of this phenomenon. Notably, phase separation was observed in native ER membranes without protein overexpression, indicating that CE accumulation alone is sufficient to remodel ER membrane organization.

### CE microdomains recruit and co-confine the INSIG/SCAP/SREBP machinery

We next asked whether CE-rich microdomains selectively organize cholesterol-regulatory proteins. Candidate lipid metabolic and ER proteins were tagged with GFP, co-expressed with mCherry-SOAT1, and their enrichment within SOAT1-positive microdomains was quantified using a microdomain enrichment index (MEI; Fig. 3A). These observations were independently validated in GERVs generated from the same conditions, confirming that protein partitioning reflected intrinsic membrane organization rather than imaging artifacts (Fig. 3D-E, S3B-C).

**Figure 3.**
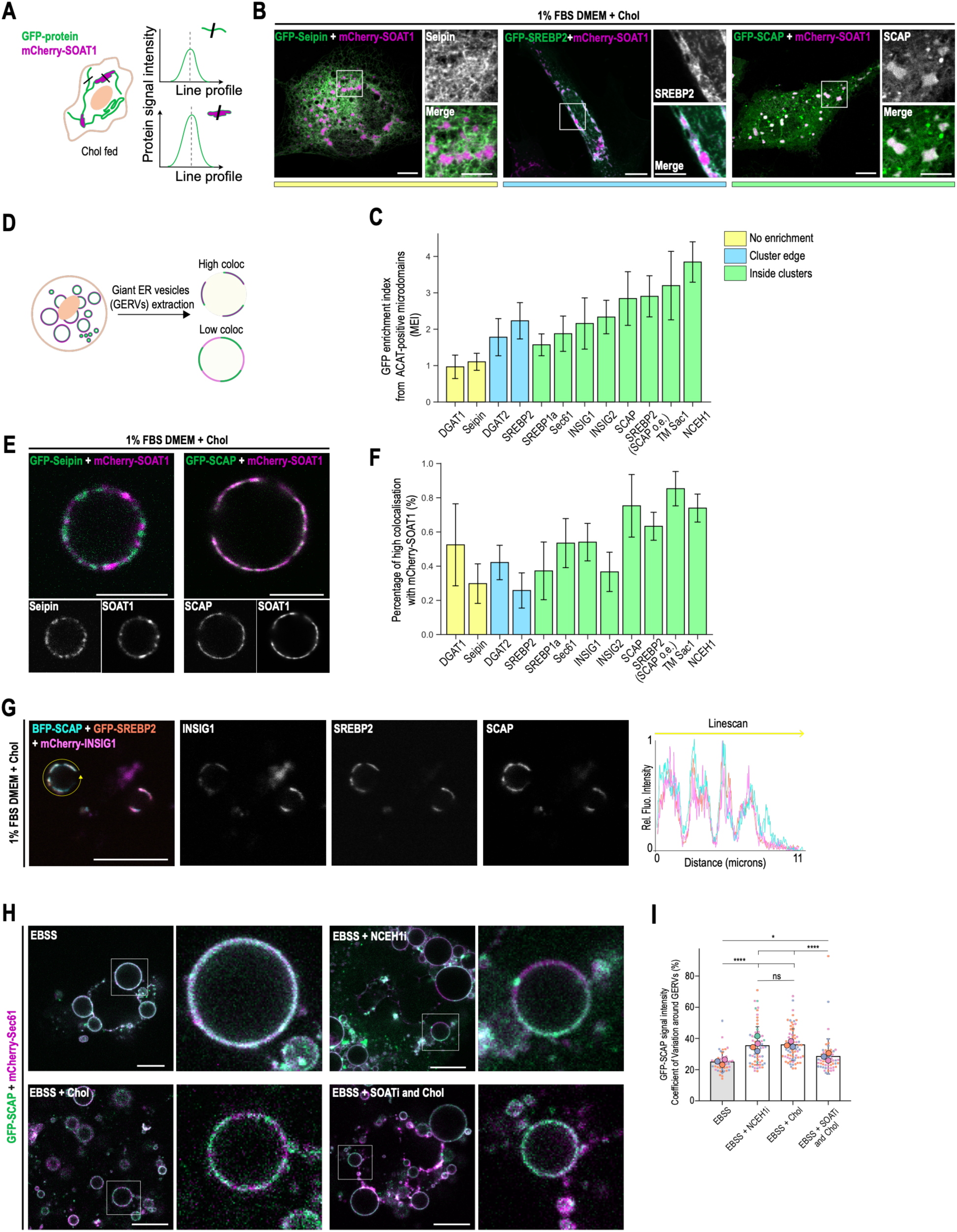
ER protein partitioning is affected by the presence of CE. **(A)** Representation of the method used to quantify the microdomain enrichment index (MEI). The intensity of the GFP signal was measured inside and outside SOAT1-positive microdomains. The MEI is the ratio of the signal from inside clusters to that from the rest of the ER. **(B)** Merged images of Huh7 cells overexpressing mCherry-SOAT1 (in magenta) with GFP-Seipin (left), GFP-SREBP2 (middle) or GFP-SCAP (right) (in green). Cells were grown in DMEM 1% FBS overnight, then fed with cholesterol at 200 μM for 2 hours. The color code (yellow, blue, and green) represents the 3 phenotypes explained in (C). **(C)** Quantification of the MEI for each GFP-tagged protein. The color code was attributed based on whether the protein is poorly/not enriched, enriched around the edges, or enriched inside the microdomains. Data collected from 3 biological replicates (n ranging from 39 to 92). **(D)** Representation of the classification applied in (E-F) to GERVs. High colocalization between GFP-tagged proteins and mCherry-SOAT1 was based on a threshold applied on peak overlap and Mander coefficients M1 + M2. **(E)** Merged images of GERVs from Huh7 cells overexpressing mCherry-SOAT1 (in magenta) and GFP-Seipin (left) or GFP-SCAP (right) (in green). Cells were grown in DMEM supplemented with 1% FBS overnight, then fed with 200 μM cholesterol for 2 hours prior to GERV extraction. **(F)** Quantification of the frequency of high colocalization signals from GERVs between SOAT1 and GFP-tagged proteins. Data collected from 3 biological replicates (n ranging from 57 to 92). **(G)** Merged image of Huh7 cells overexpressing BFP-SCAP (in cyan), GFP-SREBP2 (in orange), and mCherry-INSIG1 (in pink) grown in DMEM 1% FBS and loaded with cholesterol at 200 μM for 2 hours. The yellow arrow represents the linescan. **(H)** Merged images of swollen Huh7 cells overexpressing GFP-SCAP (in green) and mCherry-Sec61 (in magenta). Cells were starved overnight without treatment (EBSS, top left), or with NCEH1 inhibitor (EBSS + NCEH1i, top right), fed with cholesterol at 200 μM for 2 hours without any drug (EBSS + Chol, bottom left) or in the presence of SOAT inhibitor (EBSS + SOATi and Chol, bottom right). **(I)** Quantification of the coefficient of variation of GFP-SCAP signal from conditions shown in (H). Data collected from 3-4 biological replicates (n=48 EBSS; n=68 EBSS+NCEH1i; n=76 EBSS+Chol; n=59 EBSS+SOATi+Chol). Significant differences between groups were determined using a Kruskal-Wallis test followed by post hoc pairwise Mann-Whitney U tests with Bonferroni correction. Scale bars in (B), (G) and (H) represent 10 µm; scale bars in (E) and insets in (B) show 5 µm.

Proteins involved in LD biogenesis displayed distinct behaviors. DGAT1 and seipin were excluded from microdomains or showed no preferential enrichment (MEI < 1), whereas SEC61β exhibited modest enrichment (MEI ≈ 1.5). Interestingly, DGAT2 localized preferentially at microdomain boundaries (Fig. S3A), raising the possibility that boundary localization facilitates CE transfer from ER microdomains toward nascent LDs through TAG synthesis.

In contrast, the enzymes directly controlling CE metabolism were highly enriched within the same domains. NCEH1 strongly co-segregated with SOAT1, indicating that CE synthesis and hydrolysis are spatially organized within a common membrane compartment. Strikingly, the core SREBP regulatory machinery was also selectively recruited: SCAP, INSIG1, and INSIG2 were strongly enriched in microdomains and displayed extensive lateral co-segregation with SOAT1 in GERVs (Fig. 3C, F). SREBP1a similarly partitioned into microdomains. SREBP2, however, preferentially localized at microdomain boundaries. Co-expression with SCAP efficiently recruited SREBP2 into SOAT1-positive domains (Fig. 3C, F), suggesting that SCAP functions as the cholesterol-responsive adaptor that mediates SREBP2 recruitment into CE-rich membrane environments. Consistently, simultaneous expression of INSIG, SCAP, and SREBP2 resulted in near-complete lateral co-segregation of all three components (Fig. 3G). Together, these data reveal selective protein partitioning into CE-rich microdomains, with a particularly strong enrichment of the SREBP regulatory machinery.

To determine whether CE synthesis dynamically controls SCAP organization, we generated enlarged ER vesicles by controlled cell swelling and monitored SCAP and SEC61β distribution at 37°C across the four CE-manipulation conditions. Under nutrient-deprived conditions, SCAP and SEC61β displayed homogeneous co-distribution throughout the ER membrane. Cholesterol loading induced pronounced lateral segregation of SCAP into discrete domains separated from SEC61β-enriched regions. Importantly, SOAT1 inhibition prevented this redistribution despite cholesterol availability, whereas NCEH1 inhibition induced SCAP segregation in the absence of exogenous cholesterol (Fig. 3H-I). Thus, CE synthesis, rather than cholesterol loading itself or global membrane remodeling, is the determinant of SCAP lateral organization.

Consistent with this mechanism, overnight LDL loading also induced SCAP segregation in non-starved cells maintained in DMEM containing 1% FBS (Fig. S4A-D), demonstrating that CE-dependent SCAP organization occurs during physiological cholesterol uptake pathways. These findings identify CE-rich ER microdomains as a spatial platform for assembly of the cholesterol-sensing machinery.

### CE microdomains arrest SCAP within mechanically rigid and diffusion-reduced membrane regions

Phase-separated membrane domains are characterized by reduced molecular mobility and increased mechanical rigidity^31^. We asked whether CE-rich ER microdomains impose these biophysical properties on resident proteins. FRAP measurements of GFP-SOAT1 in living cells revealed substantially slower fluorescence recovery within microdomains than in the surrounding ER membrane, indicating that association with CE-rich domains strongly restricts lateral diffusion (Fig. 4A-B).

**Figure 4.**
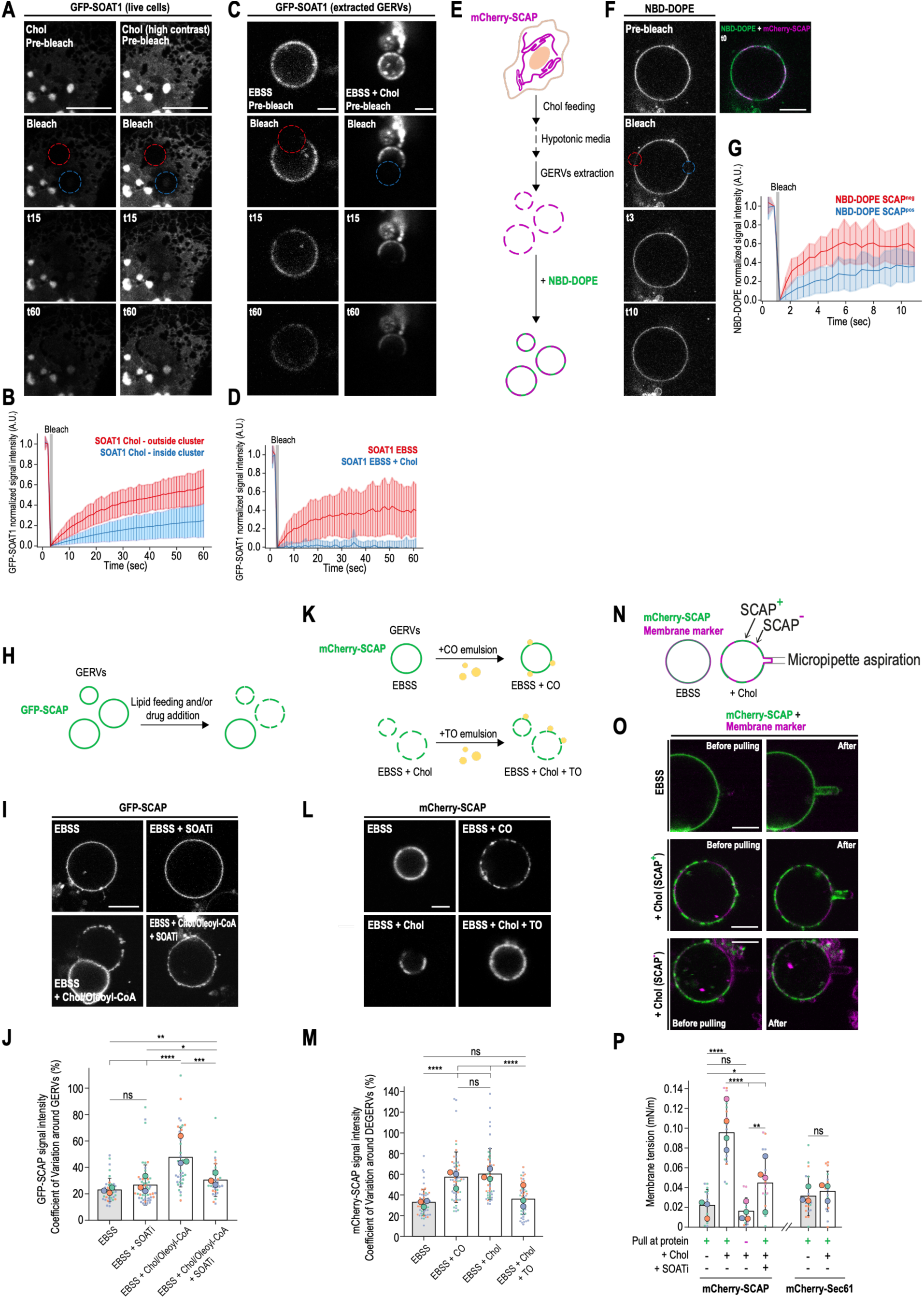
CE microdomains are diffusion-restricted membrane regions. **(A)** Time-lapse images of Huh7 cells overexpressing GFP-SOAT1, starved in EBSS overnight and fed with cholesterol at 200 µM for 2 hours. Cells were photobleached either inside (blue circle) or outside (red circle) microdomains, and fluorescence was monitored for 1 minute. **(B)** Quantification of the FRAP experiment shown in (A), with the blue line showing the recovery inside microdomains and the red line outside of them. Data collected from 3 biological replicates (n=16 outside cluster; n=16 inside cluster). **(C)** Time-lapse images of GERVs from Huh7 overexpressing GFP-SOAT1, starved in EBSS overnight alone (EBSS, left panels) or fed with cholesterol at 200 µM for 2 hours (EBSS + Chol, right panels). **(D)** Quantification of the FRAP experiment shown in (C), with the blue line showing the recovery from the EBSS + Chol condition and the red line the recovery from EBSS alone. Data collected from 3 biological replicates (n=16 EBSS; n=11 EBSS + Chol). **(E)** Representation of the main steps performed to obtain GERVs labeled with NBD-DOPE. **(F)** Time-lapse images of GERVs from Huh7 cells overexpressing mCherry-SCAP and labeled with NBD-DOPE. Cells were starved overnight and fed with cholesterol at 200 µM for 2 hours. NBD signal was photobleached inside (blue circle) or outside SCAP-positive regions (red circle), and the fluorescence was monitored for 10 seconds. **(G)** Quantification of the FRAP experiment shown in (F), with the blue line showing the recovery inside SCAP-positive domains and the red line inside SCAP-negative regions. Data collected from 3 biological replicates (n=13 SCAP^neg^; n=17 SCAP^pos^) **(H)** Representation of CE synthesis induction directly on GERVs. GERVs were extracted from Huh7 cells starved in EBSS overnight. SOAT inhibitor and/or cholesterol + oleoyl-CoA were added directly to GERVs at 37°C for 30 minutes before imaging. **(I)** Images of GERVs extracted from Huh7 cells overexpressing GFP-SCAP. GERVs from EBSS condition were used as a control (EBSS, top left), treated with SOAT inhibitor (EBSS + SOATi, top right), treated with cholesterol (100 µM) + oleoyl-CoA (50 µM) (EBSS + Chol/Oleoyl-CoA, bottom left), or with SOAT inhibitor + cholesterol + oleoyl-CoA (EBSS + SOATi + Chol/Oleoyl-CoA, bottom right). **(J)** Quantification of the coefficient of variation of GFP-SCAP signal from conditions shown in (I). Data collected from 3 biological replicates (n=40 EBSS; n=53 EBSS + SOATi; n=38 EBSS + Chol/Oleoyl-CoA; n=34 EBSS + SOATi + Chol/Oleoyl-CoA). Significant differences between groups were determined using a Kruskal-Wallis test followed by post hoc pairwise Mann-Whitney U tests with Bonferroni correction. **(K)** Representation of droplet-embedded GERVs (DEGERVs) experiment. An emulsion of cholesteryl oleate (CO) was added to GERVs extracted from EBSS alone (top panel), or an emulsion of triolein (TO) was added to GERVs extracted from EBSS + cholesterol 200 µM for 2 hours (bottom panel). **(L)** Images of GERVs extracted from Huh7 overexpressing mCherry-SCAP. GERVs extracted from EBSS (top left) were mixed with CO emulsion (top right), or GERVs extracted from EBSS + cholesterol (bottom left) were mixed with TO emulsion (bottom right). **(M)** Quantification of the coefficient of variation of mCherry-SCAP signal from conditions shown in (L). Data collected from 3 biological replicates (n=58 EBSS; n=53 EBSS + CO; n=48 EBSS + Chol; n=49 EBSS+Chol+TO). Significant differences between groups were determined using a Kruskal-Wallis test followed by post hoc pairwise Mann-Whitney U tests with Bonferroni correction. **(N)** Representation of micromanipulation of GERVs. **(O)** Merged images of GERVs from Huh7 cells overexpressing mCherry-SCAP (in green) and labeled with Laurdan (in magenta). GERVs were extracted from EBSS alone (top panel), EBSS + cholesterol and pulled at SCAP-positive regions (middle panel) or EBSS + cholesterol and pulled at SCAP-negative regions (bottom panel). Left panels show before pulling, and right panels show the GERVs during pulling. **(P)** Quantification of membrane tension from GERVs shown in (O) and more. Data collected from 3-4 biological replicates (from left to right: n=15-18-15-18-23-16). Significant differences between groups were determined using a Kruskal-Wallis test followed by post hoc pairwise Mann-Whitney U tests with Bonferroni correction. Scale bars in (A), (F) and (I) represent 10 µm, in (O) 5 µm, in (B) and (L) 2 µm.

To isolate the contribution of the membrane itself, we performed FRAP on GERVs. In GERVs derived from nutrient-deprived cells, GFP-SOAT1 recovered rapidly after photobleaching, consistent with unrestricted diffusion. In contrast, GERVs isolated from cholesterol-loaded cells exhibited minimal recovery when GFP-SOAT1 was bleached within domain regions, indicating diffusional reduction (Fig. 4C-D). GFP-SCAP displayed the same behavior (Fig. S5A-B). Similarly, the lipid probes NBD-DOPE and NBD-DPPE exhibited markedly reduced mobility within SCAP-positive regions compared with the surrounding membrane (Fig. 4E-G and Fig. S5C-D). Raising the temperature restored mobility (Fig. S5E-F), consistent with a reversible membrane domain phase transition. Furthermore, NBD-labeled cholesteryl ester preferentially partitioned into SCAP-containing domains following cholesterol loading (Fig. S5G-H), supporting the conclusion that these structures are enriched in CE.

We next asked whether CE synthesis is sufficient to generate diffusion-restricted domains in real time. GERVs isolated from starved cells, in which SOAT1 and SCAP were initially uniformly distributed, were supplemented with cholesterol, oleoyl-CoA, and ATP to drive endogenous cholesteryl oleate synthesis. This biochemical reconstitution triggered progressive formation of membrane domains accompanied by redistribution of SCAP into the newly formed regions (Fig. 4H-J, Fig. S5I, Videos S6). Inhibition of SOAT1 completely prevented domain formation under identical conditions (Fig. 4H-J, Fig. S5I, Video S7).

The direct addition of cholesteryl oleate aLDs to homogeneous GERVs induced domain formation and protein confinement, whereas TAG addition dissolved pre-existing domains (Fig. 4K-M). Thus, CE synthesis is both necessary and sufficient to reversibly generate diffusion-restricted membrane domains, dissolved by TAG.

We next examined whether CE-rich domains also exhibit distinct mechanical properties. Micropipette aspiration showed that GERVs isolated from starved cells behaved as soft, fluid membranes, whereas pulling on domain-containing regions in GERVs from cholesterol-loaded cells required substantially higher suction pressures, indicating increased membrane rigidity (Fig. 4N-P). This stiffening was markedly reduced by SOAT1 inhibition. In contrast, pulling on SEC61β-positive regions showed no difference between conditions (Fig. 4Q), demonstrating that mechanical stiffening is confined to CE-rich domains rather than representing a global change in ER membrane mechanics. These measurements establish that CE-driven phase separation generates membrane regions that are simultaneously mechanically more rigid and diffusion restricted.

These observations suggested that CE-rich domains may facilitate assembly of the INSIG/SCAP-SREBP complex by increasing local protein concentration while limiting diffusional escape. To evaluate this possibility, we modeled two-dimensional diffusion on the ER membrane using experimentally measured diffusion coefficients and conservative estimates of membrane area, cholesterol abundance, and SCAP and INSIG copy numbers (Supplementary Text). Cholesterol molecules at 5% in the ER are predicted to encounter SCAP within less than 1 ms, whereas SCAP-INSIG encounters occurred within approximately 30s. These estimations indicate that molecular encounter itself is not rate limiting. Therefore, the limiting step is not encountering but the productive complex formation, which requires cholesterol-bound SCAP to adopt the appropriate conformation, expose its MELADL motif, and engage INSIG in the correct membrane geometry. By co-confining SCAP, INSIG, and cholesterol within diffusion-restricted membrane domains, CE phase separation may increase local reactant concentration and reduce diffusional escape, thereby accelerating productive complex assembly.

### Cholesterol ester levels directly regulate SCAP retention and SREBP silencing

To determine whether CE microdomain formation regulates SCAP retention and SREBP silencing, we examined SCAP localization to the Golgi apparatus and SREBP2 nuclear translocation under the four conditions described above. In cholesterol-replete cells, GFP-SCAP and GFP-SREBP2 were largely retained in the ER (Fig. 5A-D). Inhibition of SOAT1 promoted SCAP accumulation at the Golgi and increased nuclear SREBP2 despite excess cellular cholesterol. Conversely, inhibition of NCEH1 retained SCAP in the ER and reduced SREBP2 nuclear accumulation during nutrient deprivation, a condition that normally promotes SCAP/SREBP export (Fig. 5E-H).

**Figure 5.**
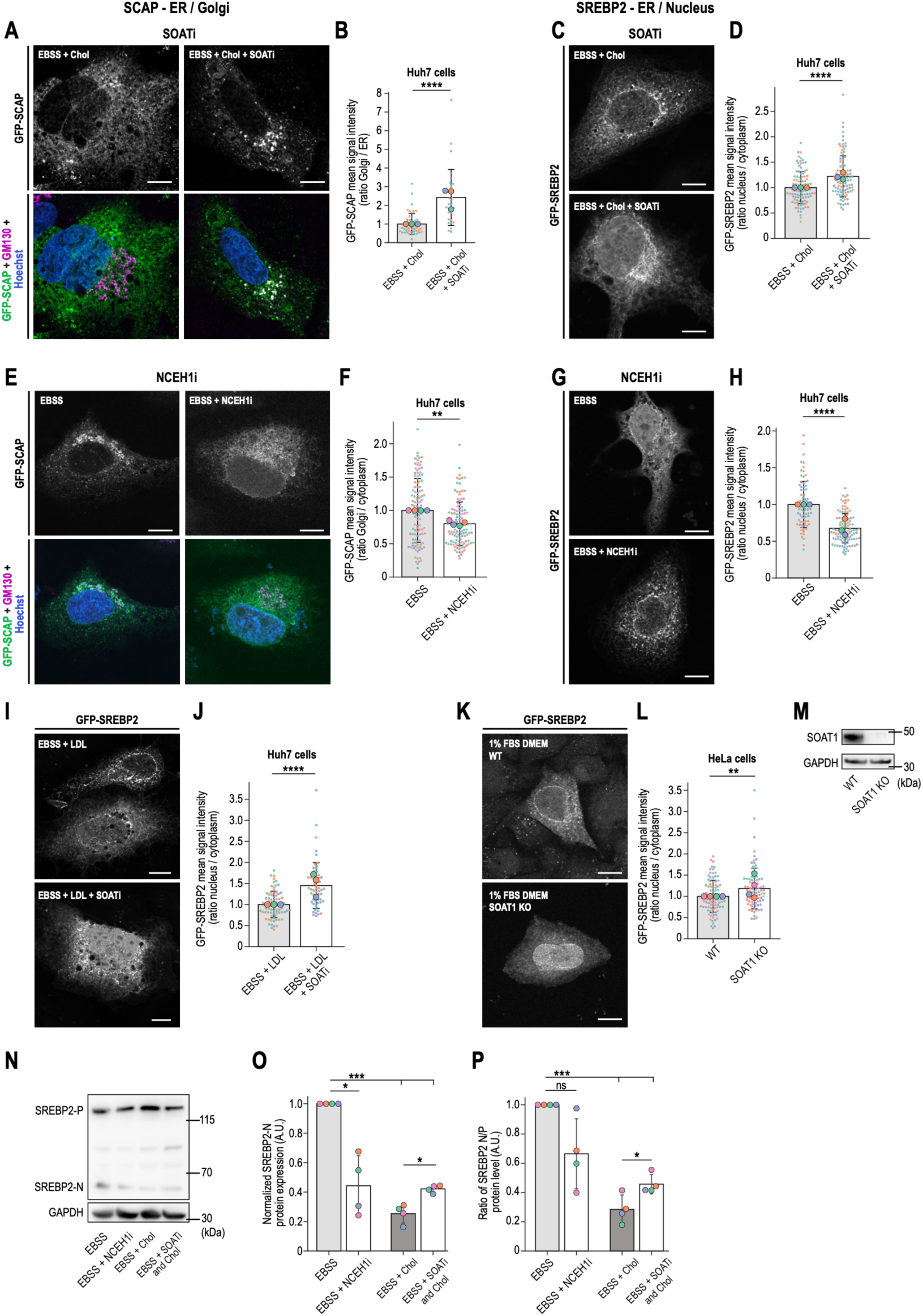
Presence of CE is required for SCAP retention and SREBP2 silencing. **(A)** Images of IF from Huh7 cells overexpressing GFP-SCAP and labeled with anti-GM130 and Hoechst. Cells were starved overnight in EBSS, then fed with 200 µM cholesterol for 2 hours, either alone (left panels) or in the presence of a SOAT inhibitor (right panels). **(B)** Quantification of GFP-SCAP signal ratio between Golgi and ER from conditions shown in (A). Data collected from 3 biological replicates (n=51 EBSS + Chol; n=31 EBSS + SOATi + Chol). A Mann-Whitney U test was performed for statistical analysis. **(C)** Images of Huh7 cells overexpressing GFP-SREBP2. Cells were starved overnight in EBSS, then fed with cholesterol at 200 µM for 2 hours, either alone (top) or with the SOAT inhibitor (bottom). **(D)** Quantification of GFP-SREBP2 signal ratio between the nucleus and ER from conditions shown in (C). Data collected from 3 biological replicates (n=101 EBSS + Chol; n=102 EBSS + SOATi + Chol). A Mann-Whitney U test was performed for statistical analysis. **(E)** Images of IF from Huh7 cells overexpressing GFP-SCAP and labeled with anti-GM130 and Hoechst. Cells were starved overnight in EBSS alone (left panels) or in the presence of the NCEH1 inhibitor (right panels). **(F)** Quantification of GFP-SCAP signal ratio between Golgi and ER from conditions shown in (E). Data collected from 4 biological replicates (n=114 EBSS; n=120 EBSS + NCEH1i). A Mann-Whitney U test was performed for statistical analysis. **(G)** Images of Huh7 cells overexpressing GFP-SREBP2. Cells were starved overnight in EBSS alone (top) or with NCEH1 inhibitor (bottom). **(H)** Quantification of GFP-SREBP2 signal ratio between the nucleus and ER from conditions shown in (G). Data collected from 3 biological replicates (n=79 EBSS; n=111 EBSS + NCEH1i). A Mann-Whitney U test was performed for statistical analysis. **(I)** Images of Huh7 cells overexpressing GFP-SREBP2. Cells were fed with LDL at 100 µg/mL in EBSS overnight alone (top) or in the presence of SOAT inhibitor (bottom). **(J)** Quantification of GFP-SREBP2 signal ratio between the nucleus and ER from conditions shown in (I). Data collected from 3 biological replicates (n=77 LDL; n=58 LDL + SOATi). A Mann-Whitney U test was performed for statistical analysis. **(K)** Images of HeLa WT and SOAT1 KO cells overexpressing GFP-SREBP2. Cells were grown in DMEM with 1% FBS overnight alone (top) or in the presence of SOAT inhibitor (bottom). **(L)** Quantification of GFP-SREBP2 signal ratio between the nucleus and ER from conditions shown in (K). Data collected from 4 biological replicates (n=94 WT; n=89 SOAT1 KO). A Mann-Whitney U test was performed for statistical analysis. **(M)** Western blot analysis of HeLa WT versus SOAT1 KO cells; stained against SOAT1 and GAPDH. **(N)** Western blot analysis of Huh7 cells, starved in EBSS overnight (first lane), in the presence of NCEH1 inhibitor (second lane), fed with cholesterol (200 µM) and 25-HC (625 nM) for 2 hours (third lane), fed with cholesterol and 25-HC in the presence of SOAT inhibitor (fourth lane); stained against SREBP2 and GAPDH. **(O-P)** Quantification of the protein level of the SREBP2 nuclear form (O) or the ratio of SREBP2 nuclear / precursor forms (P) from conditions shown in (N). Data collected from 4 biological replicates. Significant differences between groups were determined using a one-way ANOVA with Tukey’s post hoc test for multiple comparisons. Scale bars represent 10 µm.

We next tested whether these effects were conserved across cell types and modes of cholesterol delivery. In Huh7 and HeLa cells cultured in DMEM containing 1% FBS, SOAT1 inhibition similarly increased Golgi localization of SCAP and nuclear accumulation of SREBP2 (Fig. S6A-F). Overnight LDL supplementation likewise retained GFP-SREBP2 in the ER, whereas SOAT1 inhibition restored its nuclear localization. Consistent with these pharmacological data, HeLa SOAT1 knockout cells exhibited increased nuclear SREBP2 compared with wild-type cells (Fig. 5K-M).

Immunoblot analysis confirmed these effects at the endogenous level. Following nutrient deprivation and cholesterol replenishment, SOAT1 inhibition increased the cleaved nuclear form of SREBP2 (SREBP2-N), whereas NCEH1 inhibition reduced SREBP2 cleavage during nutrient deprivation alone (Fig. 5N-P). Similar changes were observed in the ratio of cleaved to precursor SREBP2, demonstrating that modulation of CE metabolism directly alters SREBP2 processing.

We then examined SREBP1a, which shares the SCAP-dependent transport pathway with SREBP2 while regulating both cholesterol and fatty acid biosynthesis^6,32,33^. Across Huh7 and HeLa cells, SOAT1 inhibition consistently increased SREBP1a nuclear localization, whereas NCEH1 inhibition reduced it under both nutrient-depleted and nutrient-replete conditions. Similar results were obtained using SOAT1 knockout and RNA silencing (Fig. S7G-O, S8). Thus, CE balance broadly regulates SCAP-dependent activation of both SREBP isoforms.

To determine whether oxysterol-mediated inhibition operates through the same mechanism, we compared the effects of SOAT1/NCEH1 inhibition with 25-HC. During nutrient deprivation, NCEH1 inhibition and 25-HC each suppressed SREBP1a activation to a similar extent, and their combination produced no further inhibition, consistent with convergence on a common pathway (Fig. S8A, B). In contrast, SOAT1 inhibition only partially restored SREBP1a activation in the presence of 25-HC, indicating that 25-HC suppresses SCAP trafficking through both CE microdomain formation and direct INSIG binding. Equivalent results were obtained in lovastatin-treated cells (Fig. S8C, D), supporting parallel contributions of these two mechanisms.

Finally, because statins activate SREBP2 by reducing cholesterol synthesis, we asked whether CE metabolism influences this response. In lovastatin-treated HuH7 and HeLa cells, GFP-SCAP accumulated at the Golgi as expected, whereas inhibition of NCEH1 increased ER retention of both SCAP and SREBP2 (Fig. S6G-L). These findings indicate that CE turnover can modulate the threshold for statin-induced SREBP activation by regulating the stability of ER CE microdomains.

Together, these findings demonstrate that disruption of CE microdomain assembly uncouples SCAP trafficking from cellular cholesterol abundance: cells containing excess cholesterol continue to activate SREBP when CE domains cannot form, whereas stabilization of CE microdomains reduces SREBP activation even under sterol-depleted conditions. These results identify ER CE accumulation as a critical determinant of SCAP retention that acts in concert with the established SCAP-INSIG interaction. Whether CE microdomains represent an obligate intermediate in physiological cholesterol sensing or cooperate with additional CE-independent retention mechanisms remains an important question for future studies.

## DISCUSSION

The INSIG/SCAP-SREBP pathway is a paradigmatic system for cholesterol homeostasis, yet how gradual changes in ER cholesterol are converted into a sharp transcriptional switch remains unresolved. Here, we identify CE phase separation as a spatial regulatory layer that links cholesterol excess to efficient SCAP retention and SREBP silencing. Whereas TAG-rich LDs form rapidly after fatty acid supply, CE-containing LDs appear with delayed kinetics after cholesterol loading^23^. Before CE is incorporated into LDs for long-term storage, it transiently reorganizes the ER membrane into microdomains enriched in SCAP and INSIG, marked by reduced protein mobility and enhanced SCAP retention.

These observations suggest that CE performs two temporally distinct functions. Following cholesterol entry into the ER, SOAT1-mediated esterification drives CE accumulation and the formation of transient microdomains that serve as templates for SCAP-INSIG assembly, thereby limiting SCAP transport to the Golgi. As TAG synthesis proceeds, CE is partitioned into LDs, where it serves as a storage reservoir for excess sterol. CE therefore acts first as a spatial regulator of cholesterol sensing and may perform additional functions in the ER that remain to be elucidated, e.g., serving as platforms for protein degradation. Only later does it serve as a storage metabolite, with the transition between these states governed by lipid flux through TAG biosynthesis.

Importantly, manipulating CE metabolism alters SCAP retention independently of bulk ER cholesterol abundance, indicating that the relevant signal is not simply the average ER cholesterol concentration. We propose that CE microdomains generate localized sterol environments that SCAP samples. Cholesterol within these domains may originate from both lateral diffusion from the surrounding ER membrane and local NCEH1-mediated CE hydrolysis, whereas SOAT1 continuously removes free cholesterol by re-esterification. SCAP is thus exposed to a dynamic local sterol pool determined by the balance of cholesterol delivery, hydrolysis, and esterification rather than by the mean ER cholesterol concentration. Such a localized metabolic circuit could explain how the pathway achieves high sensitivity despite continuous cholesterol flux.

This framework recasts the canonical ∼5 mol% ER cholesterol threshold as a metabolic set point rather than a simple equilibrium binding threshold. Cholesterol binding to the SCAP sterol-sensing domain remains essential for INSIG engagement, but our data indicate that efficient retention also depends on the membrane environment created by CE phase separation. The critical threshold may therefore correspond to the point at which local SOAT1/NCEH1 activity generates sufficient CE to drive cooperative microdomain formation. In this view, the ultrasensitive behavior of the SREBP pathway emerges from the integration of molecular recognition with cooperative membrane organization.

The enzymatic network regulating ER CE availability likely varies across tissues. SOAT1 is broadly expressed, whereas CE hydrolysis is mediated by several enzymes with distinct tissue preferences. NCEH1 is particularly abundant in macrophages and hepatocytes, while CES1, LIPE, and members of the AADAC family contribute to CE turnover in the liver, intestine, and other tissues^21,34^. This diversity may allow different cell types to tune their cholesterol-sensing threshold by adjusting the balance among CE synthesis, hydrolysis, and LD partitioning, suggesting that CE microdomains constitute a general regulatory principle whose quantitative properties are shaped by tissue-specific lipid metabolic networks.

CE dynamics also provide a mechanistic explanation for the asymmetric kinetics of SREBP regulation. Silencing is rapid because CE phase separation is cooperative, enabling swift assembly of SCAP retention platforms once a critical concentration is exceeded. Reactivation is slower because microdomain dissolution may require clearing CE from the ER membrane, a process opposed by continued SOAT1 activity and by exchange with CE stored in LDs. This kinetic buffering may help prevent inappropriate oscillations in cholesterol biosynthesis during transient fluctuations in sterol availability.

More broadly, our findings establish membrane phase separation as a mechanism for metabolic decision-making. A metabolic product, CE, transiently reorganizes the membrane environment, thereby controlling the signaling fate of its precursor, cholesterol, and coupling lipid chemistry directly to regulatory output. The SREBP switch is therefore not solely the consequence of protein-protein or protein-lipid interactions but an emergent property that arises from integrating enzymatic lipid conversion, membrane biophysics, and spatial organization within the ER. This principle suggests that other metabolic pathways may similarly exploit lipid phase transitions to translate gradual biochemical changes into biophysical codes for precise cellular responses.

## Acknowledgement

We thank the Thiam team for the helpful discussions, especially Alicia Damm and Helin Elhan, for the review of the manuscript, and for experimental assistance. French National Research Agency, ANR, ANR-24-CE11-3433-01 CHOLESTORAGE supported this work to A.R.T. A.R.T is the recipient of the Liliane Bettencourt Prize for Life Sciences® from the Bettencourt Schueller Foundation.

## Author contributions

M.Z. and A.R.T. designed the research and experimental plan with the help of M.O.; A.R.T. secured funding. M.Z. carried out most of the experiments, with the help of M.O., J.T. and S.K.; S.Y. and F.F. handled animals and assisted with *ex vivo* experiments; A.B. and M.MD. were involved in *in vitro* and ex vivo experiments. A.R.T. wrote the manuscript, reviewed by the co-authors.

## Supplementary Figures

**Figure S1.**
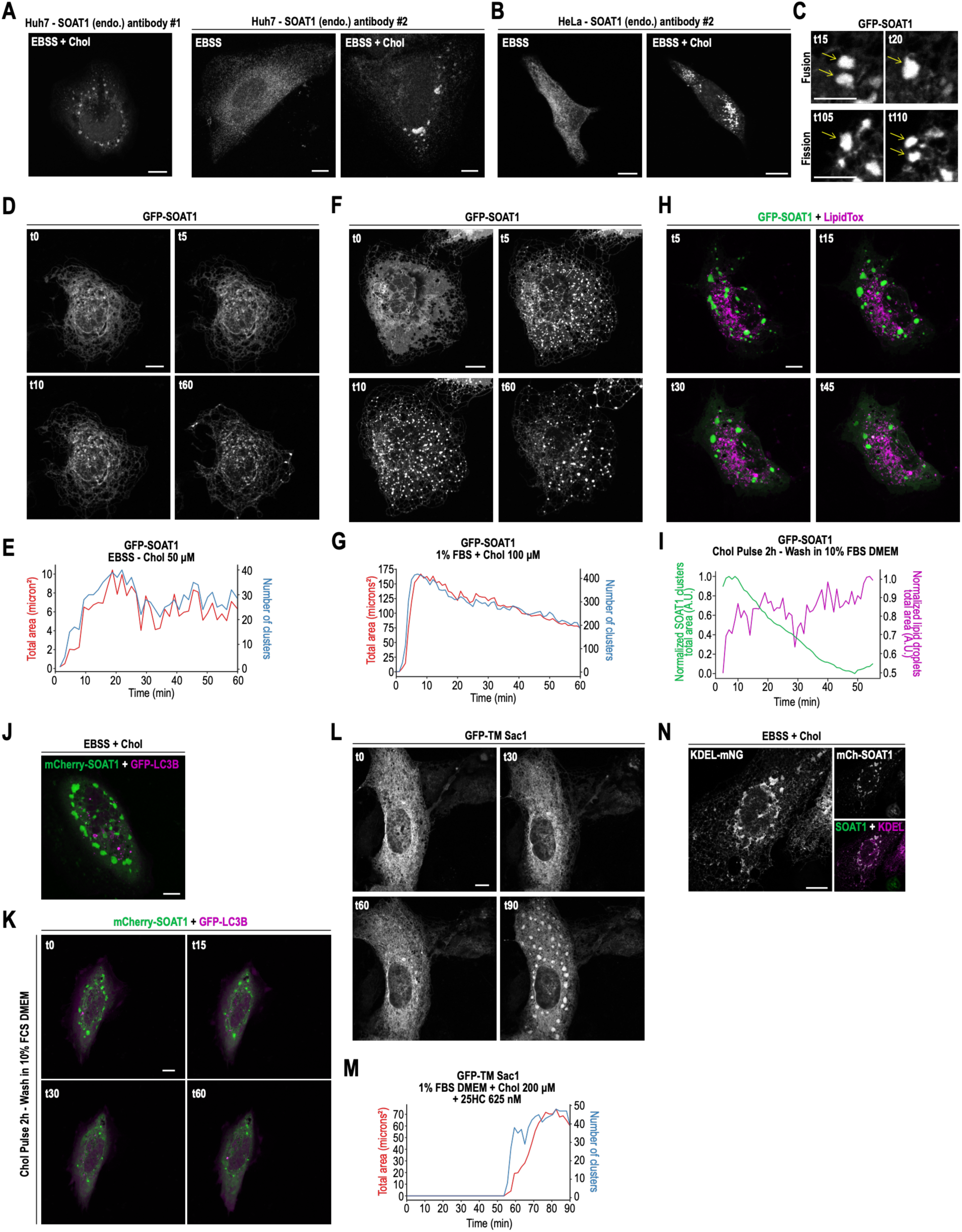
**(A)** Right. Representative immunofluorescence (IF) image of a Huh7 cell in cholesterol-loaded in EBSS + Chol (right) stained for SOAT1 (antibody #1), as in Fig.1B. Left. Representative IF image in EBSS starvation or cholesterol-loaded in EBSS + Chol, stained for SOAT1 (antibody #2). **(B)** Same experiments as in HeLa cells. **(C)** Images of microdomains in Huh7 cells overexpressing GFP-SOAT1 fed with 100 µM cholesterol. Left panels (t15-20) show a fusion event between two clusters, while right panels show a splitting event (t105-110). **(D)** Timelapse images of Huh7 cells overexpressing GFP-SOAT1. Cells were starved overnight in EBSS and fed with 50 µM cholesterol (t0 corresponds to the time of cholesterol addition). **(E)** Quantification of the total area (in red) and the number (in blue) of microdomains from the timelapse shown in (D). **(F)** Timelapse images of Huh7 cells overexpressing GFP-SOAT1. Cells were grown in DMEM + 1% FBS overnight and fed with 100 µM cholesterol (t0 corresponds to the time of cholesterol addition). **(G)** Quantification of the total area (in red) and the number (in blue) of microdomains from the timelapse shown in (F). **(H)** Timelapse images of Huh7 cells overexpressing GFP-SOAT1. Cells were starved overnight in EBSS and fed with 100 µM cholesterol for 2 hours. The medium was replaced with DMEM + 10% FBS (t0 corresponds to the time of the wash). Lipid droplets were stained using LipidTOX. **(I)** Quantification of the total area of microdomains (in green) and of lipid droplets (in magenta) from the timelapse shown in (H). **(J)** Merge image of Huh7 cells overexpressing mCherry-SOAT1 (in green) and GFP-LC3 (in magenta). Cells were starved overnight in EBSS, then fed with 100 µM cholesterol for 2 hours. **(K)** Timelapse images of the same condition as shown in (J). The medium was replaced with DMEM + 10% FBS (t0 corresponds to the time of the wash). **(L)** Timelapse images of Huh7 cells overexpressing GFP-TM Sac1. Cells were starved overnight in EBSS and fed with 50 µM cholesterol (t0 corresponds to the time of cholesterol addition). Cells were grown in DMEM + 1% FBS overnight and fed with 200 µM cholesterol + 625 nM 25-HC (t0 corresponds to the time of cholesterol addition). **(M)** Quantification of the total area (in red) and the number (in blue) of microdomains from the timelapse shown in (L). **(N)** Image of Huh7 cells overexpressing mCherry-SOAT1 (in green) and KDEL-mNeonGreen (in magenta). Scale bars represent 5 µm in panel (C) and 10 µm in the other panels.

**Figure S2.**
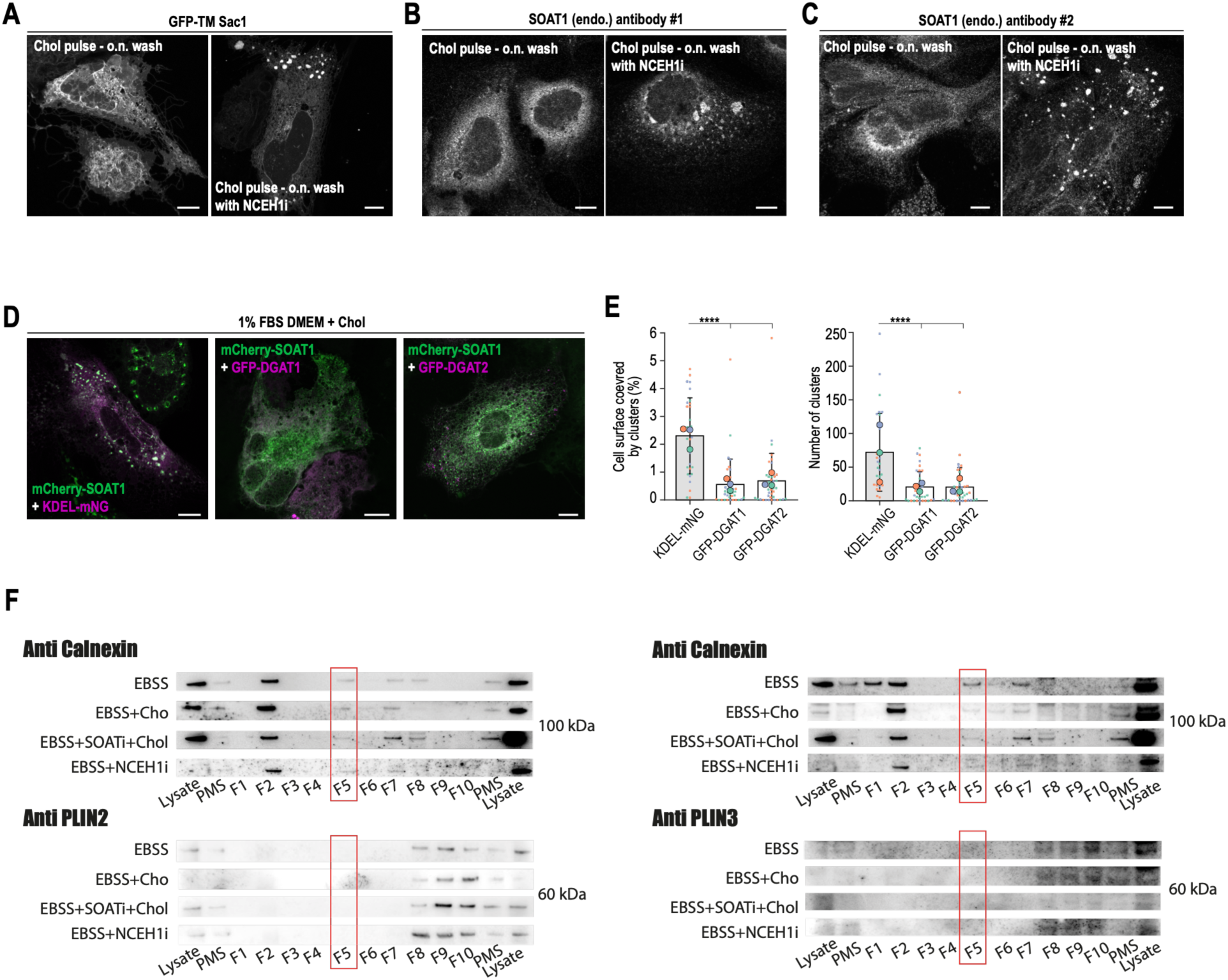
**(A)** Image of Huh7 cells overexpressing GFP-TM Sac1. Cells were fed 200 µM cholesterol for 2 hours, then the medium was replaced with fresh EBSS overnight, with or without the NCEH1 inhibitor (NCEH1i). **(B)** Images of IF experiments in Huh7 cells against SOAT1 (antibody #1) under the same conditions as shown in (A). **(C)** Images of IF experiments in Huh7 cells against SOAT1 (antibody #2) under the same conditions as shown in (A). **(D)** Merged images of Huh7 cells overexpressing mCherry-SOAT1 (in green) with KDEL-mNG (left), GFP-DGAT1 (middle), or GFP-DGAT2 (right) (in magenta). Cells were grown in DMEM + 1% FBS and fed 100 µM cholesterol for 2 hours. **(E)** Quantification of the percentage of cell surface covered by microdomains (left) and their number (right) from the conditions shown in (D). Data were collected from 3 biological replicates (n=31 KDEL; n=38 DGAT1; n=42 DGAT2). Significant differences between groups were determined using a Kruskal-Wallis test followed by post hoc pairwise Mann-Whitney U tests with Bonferroni correction. Scale bars represent 10 µm. **(F)** Example fractionation assays and blotting against calnexin and Plin2 or Plin3. Fraction 5 was analyzed. The fractionation assay was performed three times.

**Figure S3.**
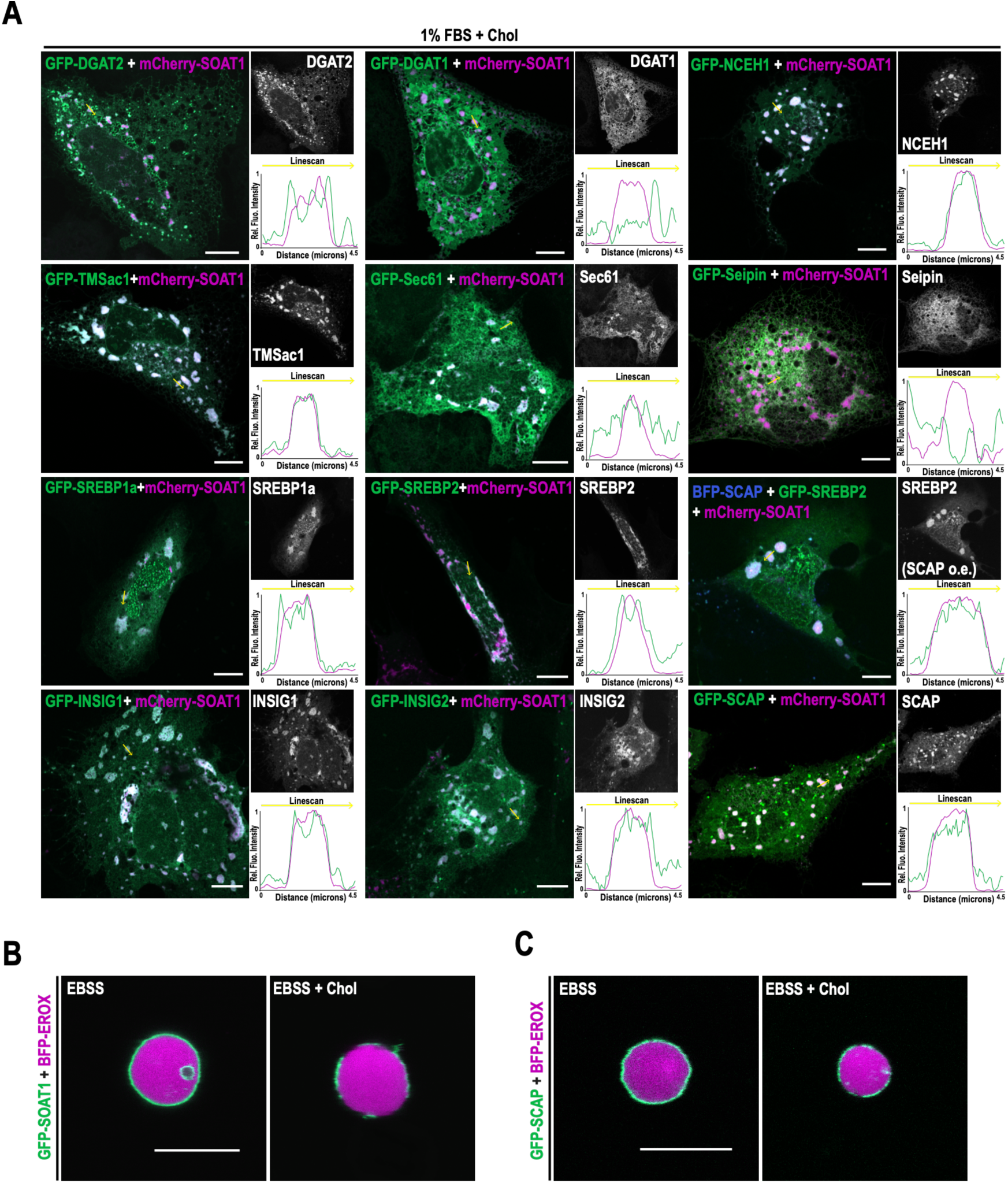
**(A)** Merged images of Huh7 cells overexpressing mCherry-SOAT1 (magenta) and a panel of GFP-tagged proteins (green). Cells were grown in DMEM + 1% FBS and fed 100 µM cholesterol for 2 hours. Yellow arrows indicate the linescans. **(B)** Merged images of GERVs extracted from Huh7 cells overexpressing GFP-SOAT1 (green) and BFP-EROX (magenta) from starved cells alone (EBSS, left) or fed cholesterol (right, EBSS + Chol). **(C)** Merged images of GERVs extracted from Huh7 cells overexpressing GFP-SCAP (green) and BFP-EROX (magenta) from starved cells alone (EBSS, left) or fed cholesterol (right, EBSS + Chol). Scale bars represent 10 µm.

**Figure S4.**
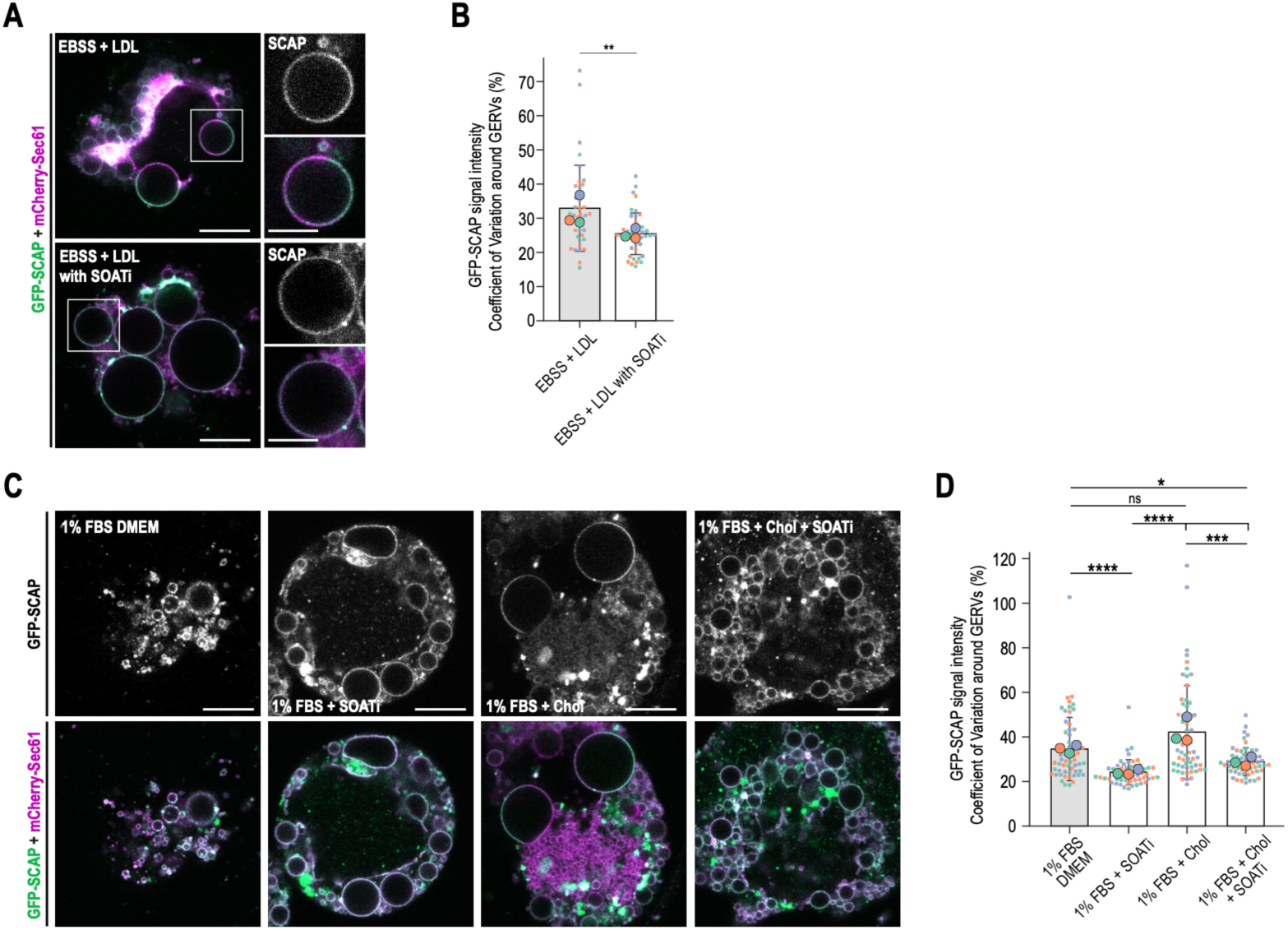
**(A)** Merged images of swollen Huh7 cells overexpressing GFP-SCAP (green) and mCherry-Sec61 (magenta). Cells were fed LDL at 100 µg/mL in EBSS overnight alone (top) or in the presence of a SOAT inhibitor (bottom). **(B)** Quantification of the coefficient of variation of GFP-SCAP signal from the conditions shown in (A). Data were collected from 3 biological replicates (n=37 LDL; n=44 LDL + SOATi). Statistical analysis was performed using a Mann-Whitney U test. **(C)** Merged images of swollen Huh7 cells overexpressing GFP-SCAP (green) and mCherry-Sec61 (magenta). Cells were grown in DMEM + 1% FBS alone (first lane), in the presence of SOATi (second lane), with the addition of cholesterol for 2 hours (third lane), or in the presence of cholesterol + SOATi (fourth lane). **(D)** Quantification of the coefficient of variation of GFP-SCAP signal from the conditions shown in (C). Data were collected from 3 biological replicates (n=60 1% FBS; n=57 1% FBS + SOATi; n=59 Chol; n=58 Chol + SOATi). Significant differences between groups were determined using a Kruskal-Wallis test followed by post hoc pairwise Mann-Whitney U tests with Bonferroni correction. Scale bars represent 10 µm; insets in (A) show 5 µm.

**Figure S5.**
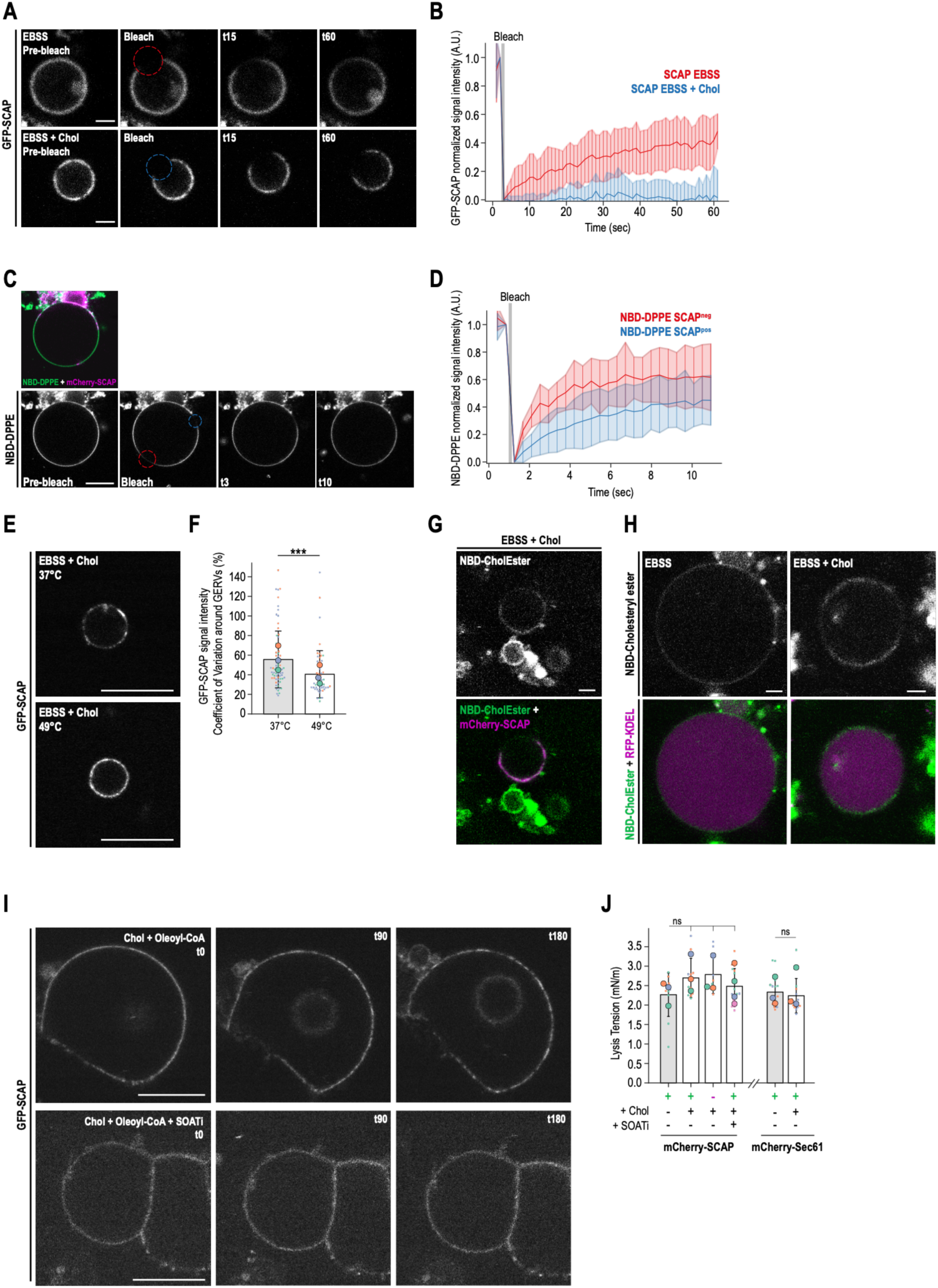
**(A)** Timelapse images of GERVs extracted from Huh7 cells overexpressing GFP-SCAP, starved overnight (EBSS, top panels) or fed with 200 µM cholesterol for 2 hours (EBSS + Chol, bottom panels). Circles indicate the photobleached areas. **(B)** Quantification of the FRAP experiment shown in (A), with the blue line showing recovery in EBSS alone and the red line in EBSS + Chol. Data were collected from 3 biological replicates (n=14 EBSS; n=10 EBSS + Chol). **(C)** Timelapse images of GERVs extracted from Huh7 cells overexpressing GFP-SCAP and labeled with NBD-DPPE. Cells were starved overnight and fed with 200 µM cholesterol for 2 hours. NBD signal was photobleached inside (blue circle) or outside SCAP-positive regions (red circle), and fluorescence was monitored for 10 seconds. **(C)** Quantification of the FRAP experiment shown in (C), with the blue line showing recovery in SCAP^pos^ domains and the red line in SCAP^neg^ regions. Data were collected from 3 biological replicates (n=14 SCAP^pos^; n=8 SCAP^neg^). **(E)** Images of GERVs extracted from Huh7 cells overexpressing GFP-SCAP and fed with cholesterol at 37°C (top) or 49°C (bottom). **(F)** Quantification of the coefficient of variation of GFP-SCAP signal from the conditions shown in (E). Data were collected from 3 biological replicates (n=72 37°C; n=51 49°C). A Mann-Whitney U test was performed for statistical analysis. **(G)** Merged images of GERVs extracted from Huh7 cells overexpressing mCherry-SCAP (in magenta) and labeled with NBD-cholesteryl ester (in green). Cells were starved overnight and fed with 200 µM cholesterol for 2 hours. **(H)** Merged images of GERVs extracted from Huh7 cells overexpressing RFP-KDEL (in magenta) and labeled with NBD-cholesteryl ester (in green). Cells were starved overnight in EBSS alone (left) or fed with 200 µM cholesterol for 2 hours (right). **(I)** Images of GERVs extracted from Huh7 cells starved in EBSS overnight. Cholesterol + oleoyl-CoA, with or without SOATi, were added to the media containing the GERVs at t0 and observed at 37°C. **(J)** Quantification of lysis tension from GERVs shown in Fig. 4O and more. Data were collected from 3-4 biological replicates (from left to right: n=11-14-10-17-19-15). Significant differences between groups were determined using a Kruskal-Wallis test followed by post hoc pairwise Mann-Whitney U tests with Bonferroni correction. Scale bars in (A), (G), and (H) represent 2 µm, and in (C), (E), and (I) 10 µm.

**Figure S6.**
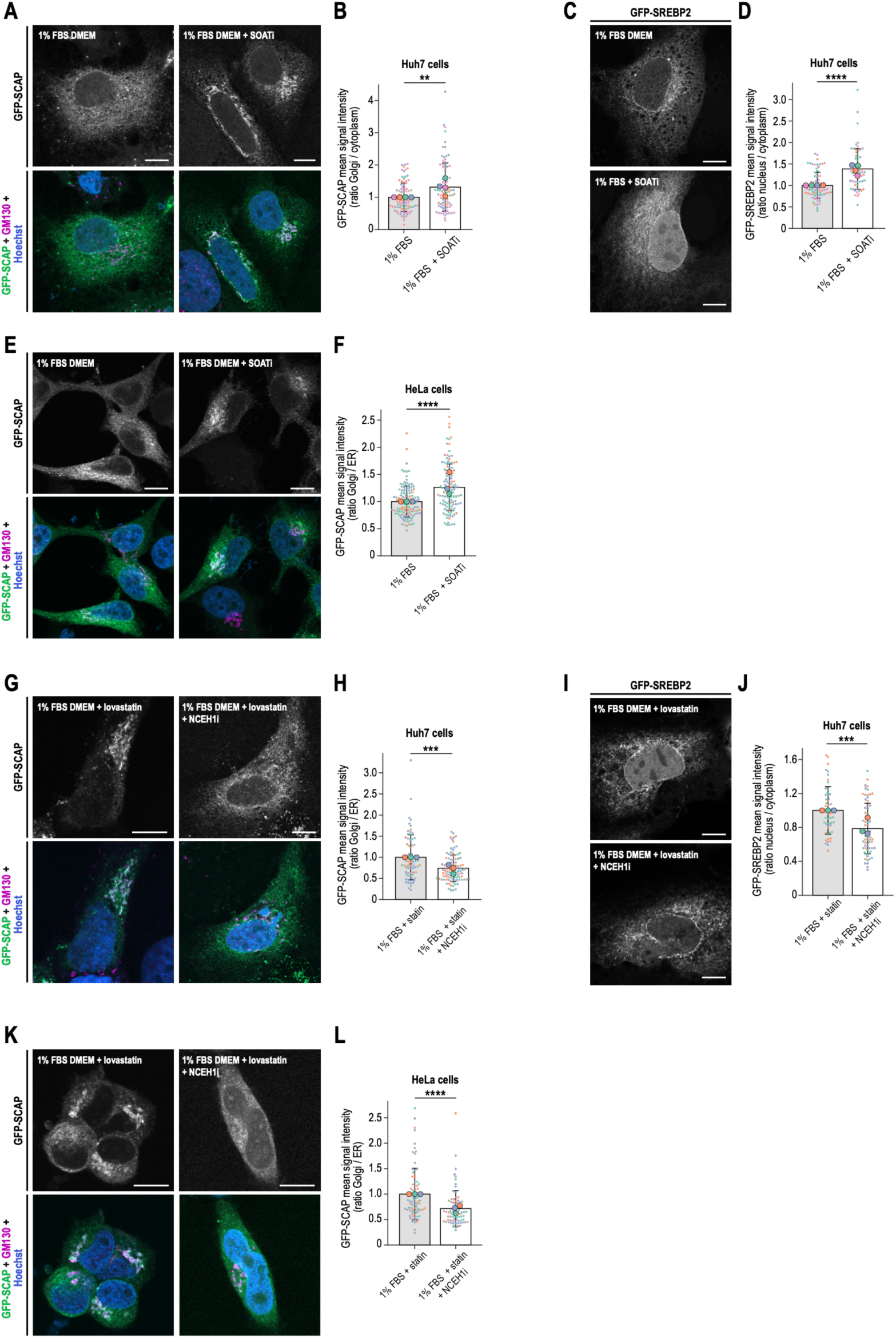
**(A)** Combined IF images of Huh7 cells overexpressing GFP-SCAP, labeled with anti-GM130 and Hoechst, grown in DMEM + 1% FBS overnight alone (left) or with SOATi (right). **(B)** GFP-SCAP signal ratio between Golgi and ER under these conditions, from 4 biological replicates (n=93 for 1% FBS; n=89 with SOATi), analyzed by Mann-Whitney U test. **(C)** IF images of Huh7 cells overexpressing GFP-SREBP2, under the same conditions as (A). **(D)** Ratio of GFP-SREBP2 signals in the nucleus versus ER, from 4 replicates (n=74 for 1% FBS; n=64 with SOATi), with Mann-Whitney U test. **(E)** IF images of HeLa cells overexpressing GFP-SCAP, labeled with anti-GM130 and Hoechst, grown in DMEM + 1% FBS overnight alone (left) or with SOATi (right). **(F)** GFP-SCAP signal ratio between Golgi and ER under these conditions, from 3 replicates (n=128 for 1% FBS; n=122 with SOATi), analyzed with Mann-Whitney U test. **(G)** IF images of Huh7 cells overexpressing GFP-SCAP, grown in DMEM + 1% FBS overnight and treated with lovastatin for 4 hours, either alone (left) or with NCEH1i (right). **(H)** GFP-SCAP ratio between Golgi and ER in these conditions, from 3 replicates (n=73 for lovastatin; n=102 with NCEH1i), with Mann-Whitney U test. **(I)** IF images of Huh7 cells overexpressing GFP-SREBP2, under the same conditions as (G). **(J)** Nuclear versus ER GFP-SREBP2 signal ratio, from 3 replicates (n=52 for lovastatin; n=65 with NCEH1i), analyzed by Mann-Whitney U test. **(K)** IF images of HeLa cells overexpressing GFP-SCAP, grown in DMEM + 1% FBS, treated with lovastatin for 4 hours alone (left) or with NCEH1i (right). **(L)** GFP-SCAP signal ratio between Golgi and ER under these conditions, from 3 replicates (n=80 for lovastatin; n=77 with NCEH1i), analyzed by Mann-Whitney U test. Scale bars = 10 µm.

**Figure S7.**
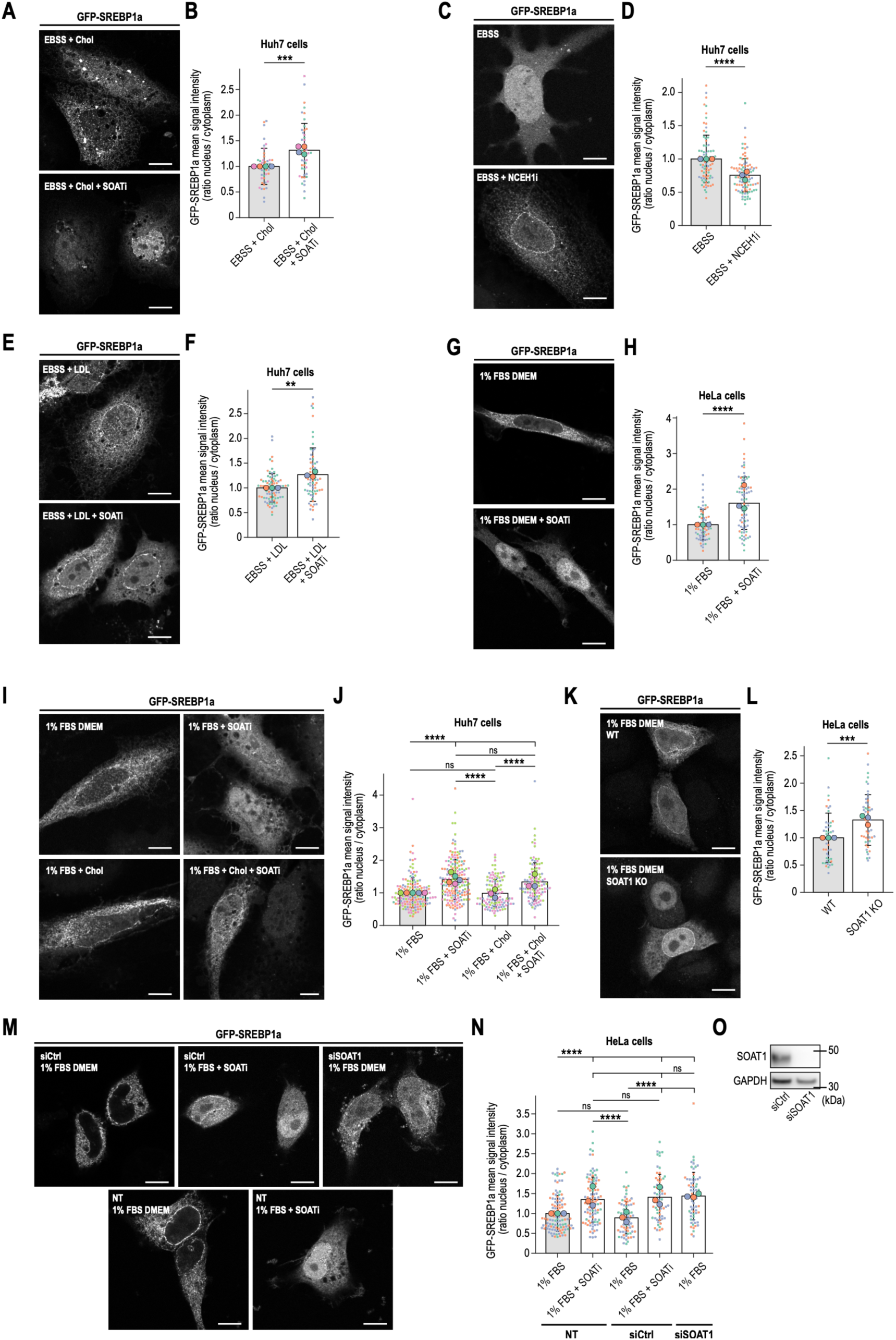
**(A)** Images of Huh7 cells overexpressing GFP-SREBP1a. Cells were starved in EBSS overnight and then fed cholesterol for 2 hours either alone (top) or with SOATi (bottom). **(B)** Quantification of the GFP-SREBP1a signal ratio between the nucleus and ER in these conditions. Data from 4 biological replicates (n=44 Chol; n=48 Chol + SOATi). Statistical analysis used Welch’s t-test. **(C)** Images of Huh7 cells overexpressing GFP-SREBP1a: cells starved overnight in EBSS alone (top) or with NCEH1i (bottom). **(D)** Quantification of the GFP-SREBP1a signal ratio between nucleus and ER in these conditions, from 3 biological replicates (n=84 EBSS; n=97 EBSS + NCEH1i), analyzed with Mann-Whitney U test. **(E)** Images of Huh7 cells overexpressing GFP-SREBP1a: cells placed in EBSS with LDL overnight alone (top) or with SOATi (bottom). **(F)** Quantification of the GFP-SREBP1a signal ratio between nucleus and ER from these conditions, with data from 3 biological replicates (n=84 LDL; n=67 LDL + SOATi), analyzed using Mann-Whitney U test. **(G)** Images of HeLa cells overexpressing GFP-SREBP1a grown in DMEM with 1% FBS alone (top) or with SOATi overnight (bottom). **(H)** Quantification of the GFP-SREBP1a signal ratio between nucleus and ER from these conditions, from 3 biological replicates (n=66 1% FBS; n=83 1% FBS + SOATi), analyzed with Mann-Whitney U test. **(I)** Images of Huh7 cells overexpressing GFP-SREBP1a: cells in EBSS overnight alone (top left) or with SOATi (top right), then fed with cholesterol for 2 hours alone (bottom left) or with SOATi (bottom right). **(J)** Quantification of the GFP-SREBP1a signal ratio between nucleus and ER, from 3-5 biological replicates (n=160 1% FBS; n=182 1% FBS + SOATi; n=119 Chol; n=114 Chol + SOATi). Significance determined by Kruskal-Wallis test and post hoc Mann-Whitney U with Bonferroni correction. **(K)** Images of HeLa WT (top) or SOAT1 KO (bottom) cells overexpressing GFP-SREBP1a grown in DMEM + 1% FBS. **(L)** Quantification of the GFP-SREBP1a signal ratio between nucleus and ER, from 3 biological replicates (n=47 WT; n=52 KO), analyzed with Mann-Whitney U test. **(M)** Images of HeLa cells overexpressing GFP-SREBP1a, transfected with control siRNA (siCtrl), SOAT1-targeted siRNA (siSOAT1), or not transfected (NT). Cells grown in DMEM 1% FBS alone or with SOATi overnight. **(N)** Quantification of GFP-SREBP1a signal ratio from these conditions, with data from 3 biological replicates (n=111 NT; n=100 NT + SOATi; n=71 siCtrl; n=64 siCtrl + SOATi; n=66 siSOAT1). Significance by Kruskal-Wallis test and post hoc Mann-Whitney U tests with Bonferroni correction. **(O)** Western blot of HeLa WT cells transfected with siCtrl or siSOAT1, stained for SOAT1 and GAPDH. Scale bars: 10 µm.

**Figure S8.**
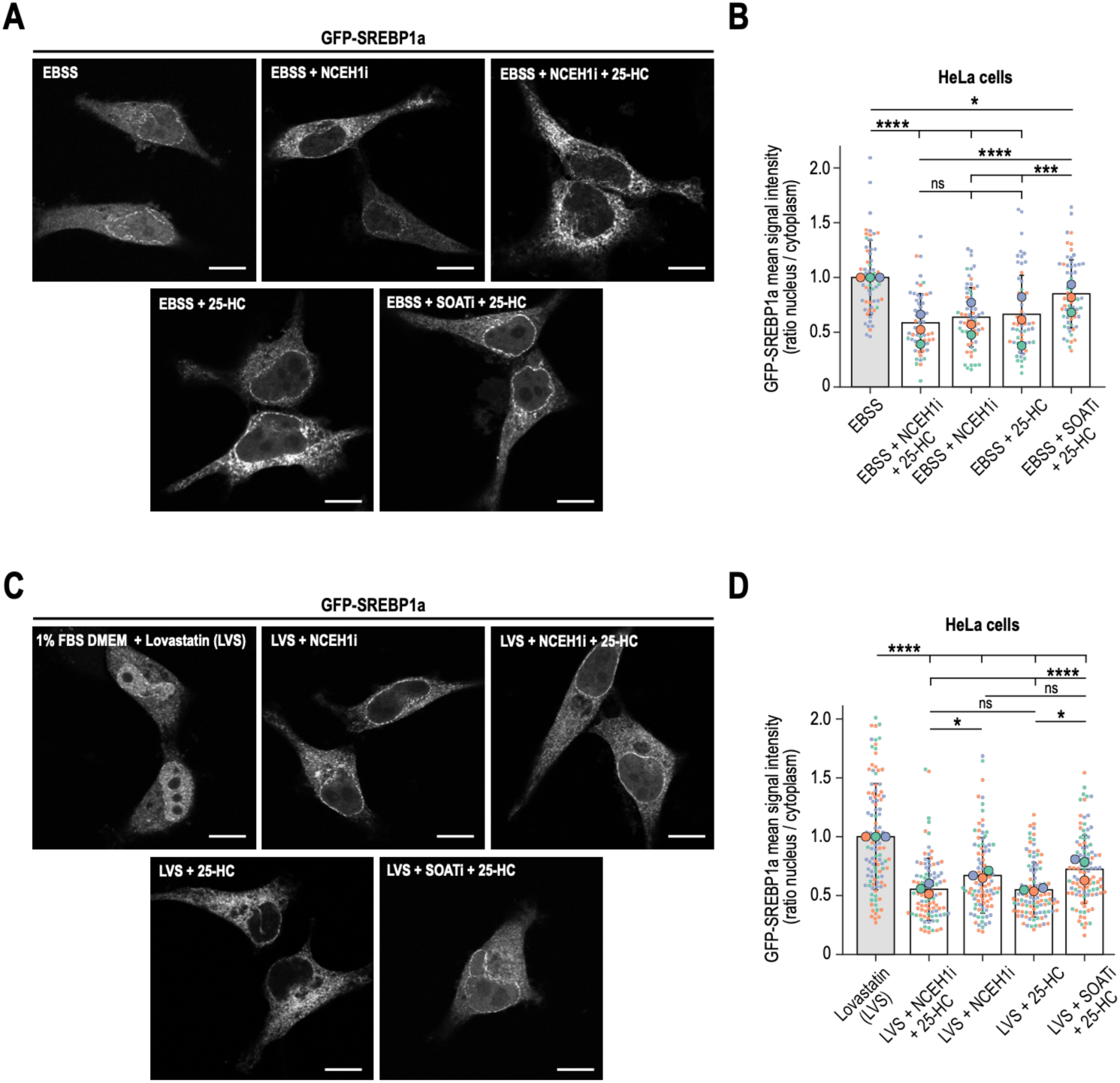
**(A)** Images of HeLa cells overexpressing GFP-SREBP1a. Cells were starved in EBSS alone (EBSS) or in the presence of NCEH1i (NCEH1i), then treated with cholesterol (200 µM) and 25-HC (625 nM) for 2 hours (EBSS + 25-HC), in the presence of NCEH1 (EBSS + NCEH1i + 25-HC) or SOATi (EBSS + SOATi + 25-HC). **(B)** Quantification of the GFP-SREBP1a signal ratio between the nucleus and ER under the conditions shown in (A). Data were collected from 3 biological replicates (n=58 EBSS; n= 61 EBSS + NCEH1i + 25-HC; n=66 EBSS + NCEH1i; n=59 EBSS + 25-HC; n=62 EBSS + SOATi + 25-HC). Significant differences between groups were determined using a Kruskal-Wallis test followed by post hoc pairwise Mann-Whitney U tests with Bonferroni correction. **(C)** Images of HeLa cells overexpressing GFP-SREBP1a. Cells were grown in DMEM + 1% FBS with lovastatin for 4 hours alone (DMEM + Lovastatin (LVS)), or in the presence of NCEH1i (LVS + NCEH1i), then treated with cholesterol (200 µM) and 25-HC (625 nM) for 2 hours (LVS + 25-HC), in the presence of NCEH1 (LVS + NCEH1i + 25-HC) or SOATi (LVS + SOATi + 25-HC). **(D)** Quantification of the GFP-SREBP1a signal ratio between the nucleus and ER under the conditions shown in (C). Data were collected from 3 biological replicates (n=108 LVS; n= 97 LVS + NCEH1i + 25-HC; n=97 LVS + NCEH1i; n=104 LVS + 25-HC; n=106 LVS + SOATi + 25-HC). Significant differences between groups were determined using a Kruskal-Wallis test followed by post hoc pairwise Mann-Whitney U tests with Bonferroni correction. Scale bars represent 10 µm.

**Figure S9.**
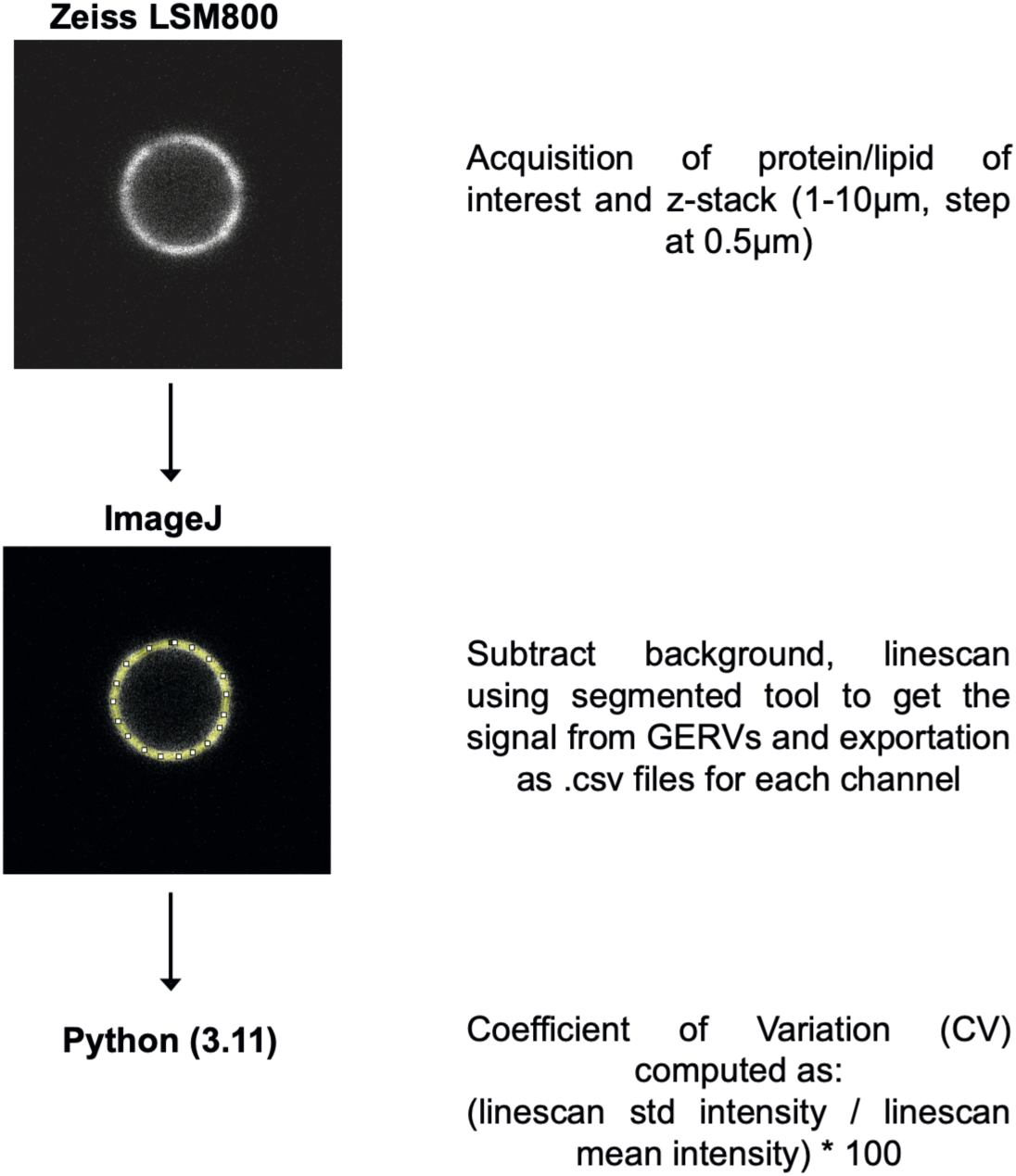
Protocol for analyzing images to calculate the coefficient of variation of protein signals on GERVs.

**Figure S10.**
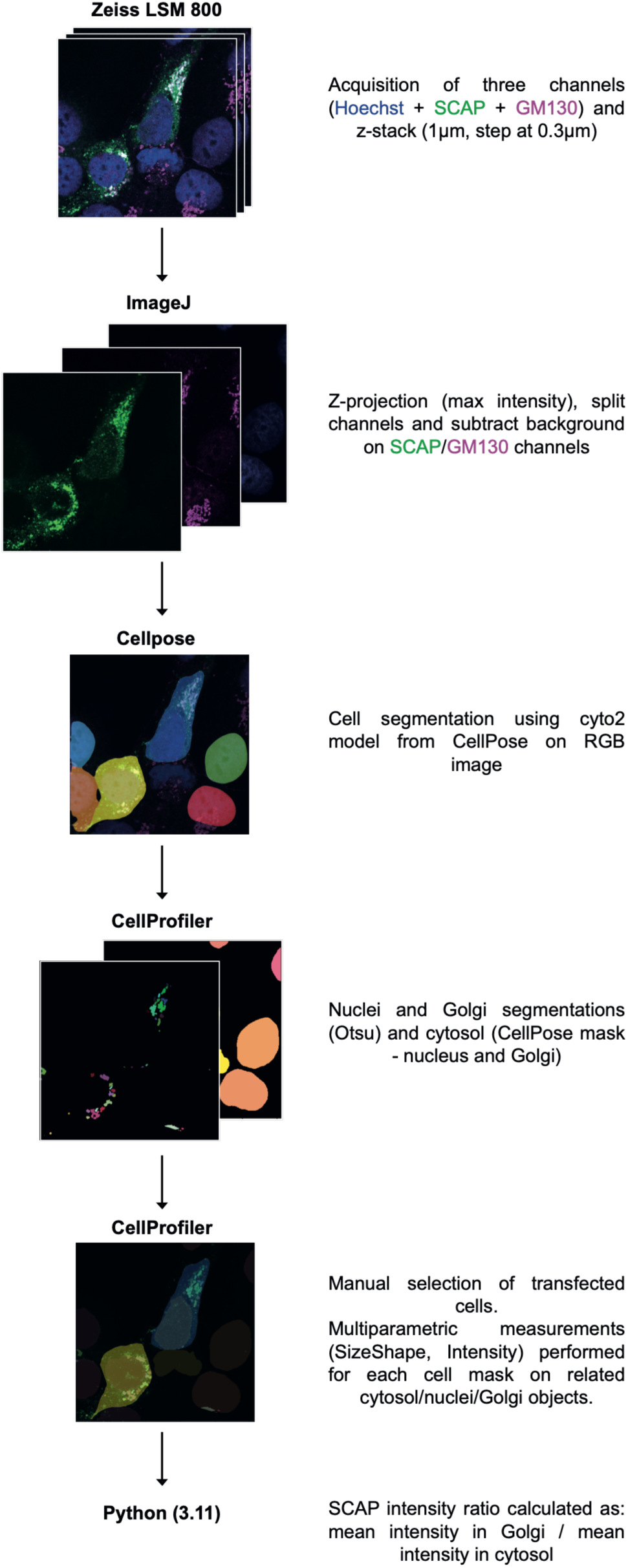
Diagram of the automated pipeline used to analyze SCAP Golgi localization under various conditions.

**Figure S11.**
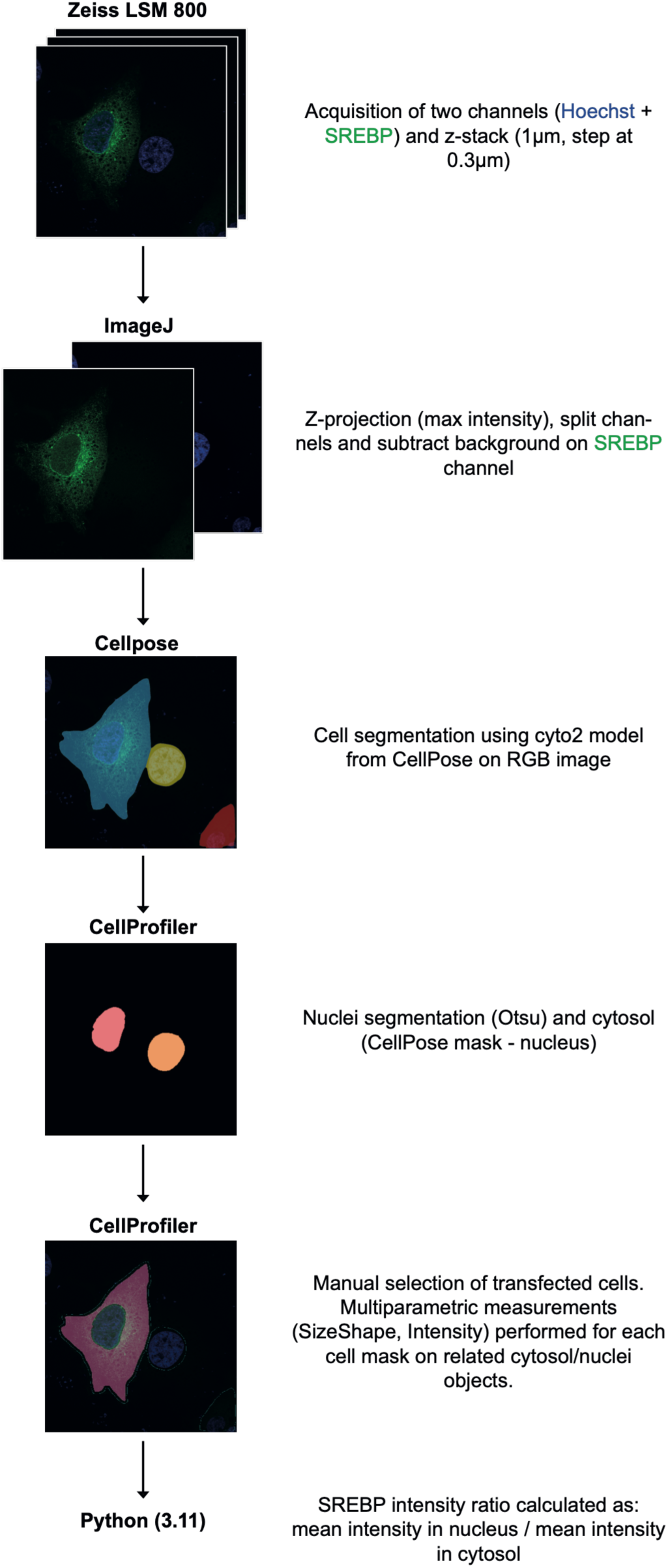
Diagram of the automated pipeline used to analyze SREBP Nuclear localization under various conditions.

---

**Movie S1:** Soat1 forms microdomain clusters following cholesterol supplementation (100µM).

**Movie S2:** Soat1 creates microdomain clusters after cholesterol is supplemented at 50µM. These clusters are fewer in number but larger than those observed with 100 µM cholesterol.

**Movie S3.** DMEM containing 1% FBS only led to a rapid burst of microdomain cluster formation.

**Movie S4:** Washing cholesterol from the culture medium and adding DMEM correlates with the disassembly of the cluster and the nucleation and growth of LDs.

**Movie S5:** Washing rapidly eliminates Soat1-GFP microdomain clusters. LC3B-mCherry does not visibly localize to the microdomain clusters.

**Movie S6:** A GERV from SCAP-GFP-transfected cells was supplemented with Oleyl-CoA and Cholesterol. Over time, SCAP becomes sequestered into microdomains.

**Movie S7:** Soat1 inhibitor was added to GERVs from SCAP-GFP-transfected cells that were subsequently supplemented with Oleyl-CoA and Cholesterol. Over time, SCAP is sequestered to microdomains.

## Material and Method

### Cell Culture

Huh-7 (gifted from Zoher Gueroui), HeLa WT and SOAT1 KO cells (gifted by Francesca Gioardano) were maintained in High Glucose (4.5 g/l) Dulbecco’s modified Eagle’s Medium (DMEM) (Dutscher) with stabilized glutamine and sodium pyruvate, supplemented with 10% fetal bovine serum (FBS) and 1% penicillin/streptomycin (GibcoBRL). Cells were cultivated at 37°C in a humidified 5% CO2 atmosphere. Mycoplasma contamination was routinely checked by PCR.

Methyl-β-cyclodextrin (#C4555) was obtained from Sigma-Aldrich, and cholesterol (Avanti, 70000 P) was sourced from Avanti. The cholesterol/methyl-β-cyclodextrin solution was prepared to contain 1 mM cholesterol as follows: methyl-β-cyclodextrin was dissolved in cell culture media and incubated with crystalline cholesterol at a molar ratio of 1:20 (cholesterol to methyl-β-cyclodextrin) for 24 hours, with agitation at 37°C. The mixture was then filtered through a 0.2 µm syringe filter and stored at −20°C until use. Unless otherwise specified, the complexed cholesterol was used at a concentration of 200 µM. Human LDL (SAE0053, Sigma-Aldrich) was added to the media at 100 µg/mL overnight. 25-Hydroxycholesterol (Cat# 700019P) was purchased from Sigma-Aldrich, solubilized in ethanol with a stock concentration of 2.5 µM, and used at a working concentration of 625 nM.

For nutrient deprivation conditions, Huh7 and HeLa cells were maintained in Earle’s balanced salt solution (EBSS) (Gibco) for 24 h. To reduce exogenous cholesterol availability in some experiments, cells were maintained in DMEM supplemented with 1% FBS for 24h.

### Chemical inhibitions

Enzymatic inhibition was achieved using Sandoz 58-035 (1 μg/mL, or 2 μg/mL in SOAT1-overexpressing cells; Sigma-Aldrich, S9318), 10 μM lovastatin (Sigma-Aldrich, 438185), 1 μM JW480 (Sigma-Aldrich, SML0792) and 5 μM of each DGAT1/2 inhibitors (Sigma-Aldrich, PZ0207, PZ0233). Lovastatin was applied for ≥ 4 h; SOAT1, NCEH1 and DGATs were inhibited overnight unless noted otherwise.

### Cell transfections and plasmids

Cells were seeded 36 h before transfections on either 35 mm glass-bottom dishes (P35G-0-20-C, MatTek Life Sciences) for live experiments or 18-mm glass coverslips (CB00180RA120MNZ0, Epredia) for fixed cells. For transfection, live cells received 1.5 µg of the indicated plasmid while fixed cells received 0.5 µg, using either jetPEI transfection reagent (10110N, PolyPlus) or X-tremeGENE 9 DNA transfection reagent (XTG9-RO, Roche).

siRNA transfections were performed using Lipofectamine RNAiMAX (Invitrogen) according to the manufacturer’s instructions. Control siRNA (D-001810-10) and SOAT1-targeting siRNAs (L-005240-00-0005) were SMARTpool ON-TARGETplus obtained from Horizon Discovery.

The plasmids GFP-SOAT1 and mCherry-SOAT1 were generous gifts from Dr. Gerald Gimpl (Johannes Gutenberg University, Mainz, Germany). EGFP-DGAT1 and EGFP-DGAT2 were kindly provided by Dr. Robert Yang (University of New South Wales, Sydney, Australia). GFP-SREBP1a was a gift from Dr. Etienne Lefai (INRAE, Auvergne, France). GFP-TM Sac1 was provided by Dr. Fabien Alpy (IGBMC, Strasbourg, France), and GFP-INSIG1 was a gift from Dr. Ron Kopito (Stanford University, California, USA). GFP-SCAP was a gift from Dr. Peter Espenshade (Johns Hopkins University School of Medicine, USA).

The plasmids mCherry-SEC61β (Addgene plasmid #121160; RRID: Addgene_121160), ERoxBFP (Addgene plasmid #68126; RRID: Addgene_68126), Seipin-mEGFP (Addgene plasmid #129719; RRID: Addgene_129719), GFP-SEC61B (Addgene plasmid #121159; RRID: Addgene_121159), and mNeonGreen-KDEL (Addgene plasmid #226568; RRID: Addgene_226568) were obtained from Addgene and were originally deposited by Dr. Christine Mayr, Dr. Erik Snapp, Dr. Elina Ikonen, Dr. Christine Mayr, and Dr. Gia Voeltz, respectively.

BFP-SCAP, NCEH1-GFP, GFP-SREBP2, mCherry-SREBP2, mCherry-INSIG1, and GFP-INSIG2 were custom-designed and synthesized by VectorBuilder (Germany).

### Thin layer chromatography

HeLa and Huh7 cells were seeded at 70% confluence and incubated overnight in complete DMEM. They were then incubated with fresh complete DMEM containing 200 µM cholesterol doped with TopFluor TMR cholesterol (Sigma Aldrich #C3045, Avanti #810385P/cholesterol) at a 99:1 molar ratio, complexed with Methyl-β-cyclodextrin, for 2 hours or overnight (16hr). Cholesterol-loaded medium was removed, and cells were washed three times with DPBS. Trypsin-EDTA (Gibco™ #15400054) was added to detach cells. Cells were recovered in complete DMEM, centrifuged, and the media was discarded. The cell pellet was washed and resuspended in DPBS. Cell counts were performed for each condition, and volumes were adjusted to obtain 107 cells/mL in each sample.

200 µL of cell suspension was added to 4 mL of methanol/chloroform (1:2), vortexed, and incubated for 30 minutes. 850 µL of water was added, the mixture was vortexed, and samples were allowed to sit at room temperature for 1 hour. The samples were vortexed again and left to separate into aqueous and organic phases. The aqueous phase was discarded, and the organic phase was dried under a stream of nitrogen. Lipids were redissolved in 50 µL of chloroform/methanol (2:1), and 20 µL were spotted on a TLC plate. Lipids were separated using a mobile phase containing Hexane:Diethyl ether:Acetic acid (80:20:1, v/v/v). Finally, TopFluor TMR was detected using a UV lamp.

### Cell fractionation for microsomal fraction analysis

Huh7 cells (∼20 million cells) were seeded at 70% confluence and incubated under the following conditions: EBSS: 18 h in EBSS; Cho: 16 h in EBSS, then 2 h in EBSS/Cho 200 µM in complex with Methyl-β-cyclodextrin; SOATi: 16 h in EBSS/Sandoz 58-035 (Sigma-Aldrich # S9318, 1 μg/mL), then 2 h in EBSS/Sandoz 58-035/Cho 200 µM in complex with Methyl-β-cyclodextrin; and NCOHi: 2 h in DMEM/1 μM JW480 (Sigma-Aldrich, SML0792), then 18 h in EBSS/1 μM JW480.

Cells were then washed three times with Dulbecco’s phosphate-buffered saline (DPBS). Following the final wash, lysis buffer (10 mM HEPES, pH 7.8; 250 mM sucrose; 25 mM KCl; 1 mM EGTA; cOmplete™ Protease Inhibitor Cocktail (Roche)) was added, and cells were detached using a cell scraper. Cells were mechanically lysed using a syringe and a 25G needle. The homogenate was centrifuged at 1,000 × g for 10 min at 4°C to remove nuclei and cellular debris. The PNS was subsequently centrifuged at 7,000 × g for 15 min at 4°C to pellet mitochondria and remaining large organelles. The supernatant was collected as the post-mitochondrial supernatant (PMS) and used as the starting material for endoplasmic reticulum (ER) microsome isolation. The PMS was mixed with an equal volume of 60% iodixanol solution to obtain a final concentration of 30% iodixanol. A discontinuous density gradient was assembled in 4-mL ultracentrifuge tubes by sequentially layering: 1 mL of PMS adjusted to 30% iodixanol at the bottom, 1 mL of 20% iodixanol, 1 mL of 10% iodixanol, and 1 mL of lysis buffer on top. Gradient tubes were balanced and centrifuged in a Beckman SW60 swinging-bucket rotor at 100,000 × g (27,300 rpm) for 2 h at 4°C. Following ultracentrifugation, fractions were carefully collected from the top of the gradient and analyzed for calnexin to identify ER-enriched fractions. Fraction n° 5 was positive for calnexin and devoid of Plin2/3 in all conditions and used for lipidomic analysis.

### Fluorescent probes

Nuclei were stained with Hoechst 33342 (0.025% v/v; 62249, Thermo Fisher). Lipid droplets were stained with BODIPY FL (0.05% v/v; D2183, Thermo Fisher) or HCS LipidTOX Deep Red (0.05% v/v; H34477, Thermo Fisher). Membranes were labeled using Laurdan (working concentration: 5 μM; D250, Thermo Fisher).

### Giant unilamellar vesicles (GUVs)

The following phospholipids were used: POPC (1-palmitoyl-2-oleoyl-sn-glycero-3-phosphocholine), DOPC (1,2-dioleoyl-sn-glycero-3-phosphocholine), and DPPC (1,2-dipalmitoyl-sn-glycero-3-phosphocholine), all from Avanti Polar Lipids. The neutral lipids used was cholesteryl oleate (CO) from Sigma-Aldrich. Membranes were fluorescently labeled with NBD-DPPE and DOPE-Cy5 (Avanti Polar Lipids), each at 1 mol%. All experiments were performed in HKM buffer (50 mM HEPES, 120 mM potassium acetate, 1 mM MgCl₂, pH 7.4, 300 ± 10 mOsm) prepared in Milli-Q water.

GUVs were generated by electroformation following Chorlay et al.^29^. Three compositions were tested (mol%):

- Mix 1: POPC/DOPC/DPPC (20/40/40)
- Mix 2: POPC/DOPC/DPPC/CO (20/40/40/30)
- Mix 3: POPC/DOPC/DPPC (20/40/40), followed by the incorporation of CO droplets after GUV formation (see below)

Lipid stocks were dissolved in chloroform, mixed at the indicated ratios, and spread onto an ITO-coated glass plate. A second ITO plate was placed on top, with a PDMS spacer defining the chamber, and the assembly was dried under vacuum for 1 h. The chamber was then filled with iso-osmotic sucrose (300 ± 10 mOsm), and a 5 V peak-to-peak, 100 Hz AC field was applied across the ITO plates for 3 h at 60°C. Vesicles were harvested with a Pasteur pipette, transferred to Eppendorf tubes, and stored at 4°C until use. Imaging was carried out at room temperature.

### Droplet-embedded vesicles (DEVs)

For Mix 3, CO droplets were prepared and fused with preformed GUVs as described in Chorlay et al ^29^. CO (3–8 mg) was first melted at 60°C and dispersed in HKM by alternating sonication and vortexing until a stable white emulsion was obtained. This emulsion was mixed with 150 μL of the GUV suspension, briefly homogenized by hand, and incubated for 5 min on a rotating tube mixer. The suspension was then loaded onto a coverslip passivated with 10% (w/w) BSA in Milli-Q water and rinsed three times with HKM.

### GERVs production and extraction

After transfection, the cultured cells were transferred into a hypotonic culture medium, DMEM: H2O (5:95% v/v) at pH 7.4, 37 °C, and 5% CO2 for 20 min to induce swelling of cells and bilayer-bounded organelles. For cell experiments on swollen cells, observations were performed at this step directly on the bottom dishes of the MatTek coverslips. For GERV experiments, cells were mechanically lysed by extensive pipetting to disrupt the cell membrane, as described in ^30^.

### Incorporation of lipids into GERVs and DEGERVs production

Fluorescent lipids (NBD-DPPE 16:0 [810144P], NBD-DOPE 18:1 [810145P], Cy5-DOPE 18:1 [810335C], and NBD-12Chol [810252P]; from Merck) stored in chloroform were dried under argon and then vacuum-desiccated for at least 4 h. The dried lipids were resuspended in HKM buffer (50 mM Hepes, 120 mM potassium acetate, and 1 mM MgCl₂ in MilliQ water; pH 7.4, 300 ± 10 mOsm)/ethanol buffer (90/10 v/v) by vortexing and sonication in three cycles. Lipids were then added to GERVs to achieve final concentrations of 11 μM (NBD-DPPE), 4.3 μM (NBD-DOPE), 7.8 μM (Cy5-DOPE), and 25 μM (NBD-12Chol), and the GERVs were incubated for 12 h at 37°C before imaging.

For experiments involving domain induction directly on GERVs, cholesterol:MβCD complex (100 μM) and oleoyl-CoA:BSA complex (870719P, Avanti Research; 50 μM) were added to GERVs and incubated for 30 min at 37°C before imaging.

To produce DEGERVs ^35^, triolein (1,2,3-tri-(9Z-octadecenoyl)-glycerol, TO [870110O], Avanti Polar Lipids, purchased from Merck) emulsions were prepared by mixing 10 µL of TO with 80 µL of HKM. For cholesteryl oleate (CO [C9253], Sigma-Aldrich), a mass of 8-12 mg was weighed and heated to 60°C to ensure a liquid phase. 80 µL of HKM was then added to the CO liquid phase. The oil/HKM mixture was vortexed (20 sec) and sonicated (20 sec), and this cycle was repeated three times until a white emulsion formed. The resulting emulsion was added to the GERVs by gentle pipetting, incubated at 50°C for 20 min, then overnight at 37°C before imaging.

### Confocal microscopy imaging

All images were acquired on a Carl ZEISS LSM800 using an oil-immersion ×63 objective with NA 1.4. EGFP, NBD, Bodipy, and Alexa 488 fluorescence was excited at 488 nm, and emission was detected between 510 and 550 nm. mCherry and Alexa 555-tagged protein fluorescence was excited at 561 nm, and emission was detected between 580 and 650 nm. Deep-red fluorescence, such as Cy5 or LipidTOX, was excited at 640 nm and detected above 650 nm. BFP and Laurdan fluorescence was excited at 405 nm and detected below 500 nm.

Unless otherwise stated, live-cell imaging, FRAP, and GERVs-related experiments were performed at 37°C and 5% CO_2_ in an incubation chamber.

### Immunofluorescence

Cells were grown on glass coverslips, fixed in 4% paraformaldehyde in PBS for 15 min, then permeabilized and blocked for 20 min at room temperature with saponin (0.05% w/v) and gelatin (0.7% w/v) in PBS. Cells were incubated with primary antibodies in the same buffer overnight at 4°C. Primary antibodies were anti-SOAT1 (antibody #1, 1:500, ab39327, Abcam; antibody #2, 1:300, ab307597, Abcam) and anti-GM130 (1:800, 610822, BD Biosciences). Cells were washed twice in PBS and incubated with secondary antibodies (AlexaFluor 488 [RRID: AB_2535792 and AB_141607] and AlexaFluor 555 [RRID: AB_2762848 and AB_162543]) and Hoechst 33342 (0.025% v/v) for 30 min at room temperature. After three washes, coverslips were mounted onto slides in ProLong Gold (P36934, Invitrogen).

### Quantification of SOAT1 clustering

Using Fiji, images underwent background subtraction and channel splitting. Cell segmentation was performed manually. Binary ER masks were generated by applying intensity thresholding (Minimum) followed by watershed to the SOAT1 signal. The number of clusters and total area were retrieved using Analyze Particles (size = 0.1-Infinity, circularity = 0.1-1). Data were exported as .csv files and further processed in Python (3.11). The total cluster area was normalized by the cell area to obtain the % of cell surface covered by clusters.

### Quantification of protein enrichment to SOAT1 clusters

The outlines of SOAT1 clusters were manually drawn in Fiji using the freehand selection tool based on the SOAT1 signal. The resulting ROIs were then used to measure GFP signal intensity. For non-cluster measurements, five ROIs of similar size were drawn in GFP-positive regions outside the clusters, sampling signal from ER tubules and sheets. Data were processed in Python (3.11), and a cluster enrichment ratio was calculated for each cell as:

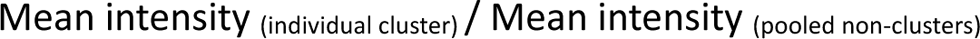

### Quantification of fluorescence signal from swollen cells or extracted GERVs

Image processing was performed using Fiji (Fig. S9). First, background subtraction was applied to all images. When multiple channels were acquired, the segmented line tool was used to manually trace the outlines of GERVs on the RGB composite. The line width was adjusted to match each GERV’s membrane signal, and lipid droplets were excluded from the analysis. Line profile intensities were measured for each channel from the corresponding ROIs and exported as .csv files. Subsequent data processing was performed in Python (3.11), where the Pearson coefficient was computed using the pandas/NumPy libraries, and the coefficient of variation (CV, %) for each GERV was calculated as:

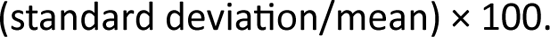

### Quantification of protein colocalization with SOAT1 on GERVs

The same method as above was used to obtain line-profile intensities for each channel (see the quantification of fluorescence signals from swollen cells or extracted GERVs). After that, .csv files were processed in Python (3.11), and intensity peaks were detected using SciPy find_peaks function. The peak overlap between channels was computed and used as a threshold (0.35) for high colocalization, in addition to the Manders metrics M1 (0.4) and M2 (0.4). GERVs satisfying these 3 thresholds were categorized as having high colocalization, and the frequency was computed for each condition.

### Animals

All animal procedures were conducted in accordance with European legislation for animal welfare (Directive 2010/63/EU). Experiments complied with French regulations (French Law 2013-118, February 6, 2013) and were approved by the relevant local ethics committees (CETEA, Comité d’Éthique en Expérimentation Animale 89, and Comité d’Éthique Charles Darwin). The approved protocol numbers were APAFIS#20687-2019051617105572 and APAFIS#53865-2025021818503991.

Mice were housed in a specific pathogen-free facility under standard conditions, including a 12 h light/12 h dark cycle, 50–70% relative humidity, and an ambient temperature of 19–21°C. Animals had ad libitum access to food and water and were maintained in enriched cages containing bedding material and gnawing sticks.

### Primary Hepatocyte Isolation and Culture

Primary hepatocytes were isolated from 14–16-week-old C57BL/6 mice by collagenase perfusion. Briefly, a 22-gauge feeding needle connected to a peristaltic pump was inserted into the inferior vena cava. Livers were first perfused with perfusion buffer (137 mM NaCl, 3.5 mM KCl, 0.2 mM Na₂HPO₄, and 10 mM HEPES, pH 7.65) at a flow rate of 6 mL/min for 3 min, followed by perfusion with collagenase buffer containing 137 mM NaCl, 3.5 mM KCl, 0.2 mM Na₂HPO₄, 10 mM HEPES, 1.87 mM CaCl₂ (pH 7.65), and 500 μg/mL Collagenase Type I (Thermo Fisher Scientific, 17018029).

Following digestion, liver lobes were transferred to washing medium (137 mM NaCl, 3.5 mM KCl, 0.2 mM Na₂HPO₄, 10 mM HEPES, and 0.5 mM EDTA, pH 7.65) and dissociated by passage through a 100 μm cell strainer. Cells were pelleted by centrifugation at 30 × g for 3 min, resuspended in fresh washing medium, and washed three times.

Viable hepatocytes were quantified by Trypan Blue exclusion using a Countess II FL Automated Cell Counter (Invitrogen) and seeded at a density of 6 × 10⁴ cells/cm². Cells were cultured in M199/G5 medium (Invitrogen, 31150030) supplemented with 100 U/mL penicillin–streptomycin (Invitrogen, 15140122), 0.1% bovine serum albumin (BSA; Sigma-Aldrich, A4503), 2% Ultroser G (Cyphergen, 15950-017), 100 nM dexamethasone (Sigma-Aldrich, D4902), 100 nM triiodothyronine (T3; Sigma-Aldrich, T6397), 1 nM insulin (Invitrogen, 03-0110-SA), and 1% L-glutamine.

### GERVs from murine primary hepatocytes

Primary hepatocytes were starved in EBSS overnight, and the cholesterol:MβCD complex was added to the media at 200 μM for 3 h before imaging. Cells were then swollen in hypotonic media containing Laurdan and LipidTOX for 20 min at 37°C. GOVs were extracted as described above and observed 12 h later. GERVs were identified by LipidTOX signal and the presence of lipid droplets. Images were processed as described above.

### Quantification of SCAP

Images underwent background subtraction, maximum-intensity z-projection, channel splitting, and high-contrast RGB conversion in Fiji (Fig. S10). Cell segmentation was performed using CellPose 2.0 (cyto2 model) on the RGB images, and the resulting cell masks, along with the split channels, were analyzed in CellProfiler (4.2.6). Transfected cells were manually selected using the EditObjectsManually module. Nuclei and Golgi objects were identified with the IdentifyPrimaryObjects module (Global, Otsu two-class thresholding). The ER/cytosol region was generated by subtracting the nuclei and Golgi objects from the cell mask. Areas and intensities for each compartment were quantified using the MeasureObjectSizeShape and MeasureObjectIntensity modules. Data were processed in Python (3.11), and the SCAP ratio for each cell was calculated as:

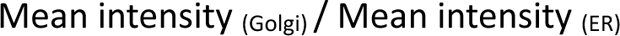

For each experiment, data were normalized by the mean value of the control condition.

### Quantification of SREBPs

Images underwent background subtraction, maximum-intensity z-projection, channel splitting, and high-contrast RGB conversion in Fiji (Fig. S11). Cell segmentation was performed using CellPose 2.0 (cyto2 model) on the RGB images, and the generated cell masks together with the split channels were analyzed in CellProfiler (4.2.6). Nuclei were identified using the IdentifyPrimaryObjects module (Global, Otsu two-class thresholding). Transfected cells were manually selected using the EditObjectsManually module. The ER/cytosol region was obtained by removing the nuclear object from the cell mask. Areas and intensities for each compartment were quantified using the MeasureObjectSizeShape and MeasureObjectIntensity modules. Data were processed in Python (3.11), and the SREBP ratio for each cell was calculated as:

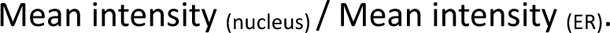

For each experiment, data were normalized by the mean value of the control condition.

### FRAP experiments

Fluorescence recovery after photobleaching (FRAP) experiments were performed by bleaching either the entire surface of SOAT1 clusters or a defined region of GERVs using the 488-nm laser line at 100% power with five iterations. Fluorescence recovery was recorded every second for 1 min in protein experiments, and every 450 ms for 12 s in lipid experiments. Image analysis was carried out in Fiji using a custom script to track objects and quantify ROI intensities over time. Data were normalized between 0 and 1 in Python (3.11), using the frame immediately before bleaching as the maximum value and the first post-bleach frame as the minimum.

### Micromanipulation of GERVs

GERVs from Huh7 cells were transfected and prepared as described above. Micromanipulation measurements were performed as previously described ^30^. Briefly, cells were swollen in diluted aqueous DMEM (5:95% v/v; Dutscher) at 37°C and 5% CO_2_ for 15 min to induce hypotonic shock. Giant organelle vesicles were then extracted by mechanically lysing the cells and placed on a BSA (10% v/v)-pre-treated coverslip. Glass pipettes (pulled from borosilicate capillaries: 1.0OD, 0.58ID, 30-0017 GC100-15b; Harvard Apparatus, Holliston, MA) were cleaned with a plasma cleaner, coated with mPEG5K-silane (Cat#JKA3037, Merck), and filled with diluted DMEM. The pipette was placed in the homemade micromanipulation setup and aligned with the microscope settings. GERVs were carefully collected using a micropipette. Aspiration pressure was measured with a pressure transducer (DP103, Validyne Engineering Corp; range: 55 kPa) attached to the pipette and controlled manually with a syringe. The output voltage of the pressure transducer was monitored with a digital voltmeter. The micromanipulators used were TransferMan 4r. Membrane tension can be calculated using Laplace’s law as follows:

(1)

With ΔP being the suction pressure, R_p_ the inner pipette radius, and R_GERV_ the radius of the GERV. Radii were obtained from CLSM images utilizing Fiji. Selected domain or non-domain parts had to be at least the pipette diameter in length.

For measurement of the ER membrane lysis tension, the initial suction pressure (πi), corresponding to the minimum pressure required to draw a membrane tongue into the micropipette, and the final suction pressure (π_end_), corresponding to membrane rupture, were recorded. The suction pressure was then ramped at a constant rate (approximately 10 mbar min⁻¹) until membrane failure occurred through pore opening. The lysis tension was calculated using Equation (1), where ΔP = π_end_ − π_i_ is the applied pressure difference. A representative pressure-loading sequence used to determine the lysis tension is shown in Fig. 4.

### Ultracentrifugation for ER fraction purification

Approximately 20 million Huh7 cells were subjected to four different treatment conditions. Following treatment, cells were washed with DPBS, scraped with a cell lifter, centrifuged, and lysed in a hypotonic buffer containing 100 mM HEPES (pH 7.8), 10 mM EGTA, 250 mM KCl, plus protease and phosphatase inhibitors (Roche). They were incubated for 20 minutes, then mechanically lysed by passing through a 25G needle 10 times. The lysate was centrifuged at 1,000 × g for 10 minutes to pellet nuclei, then at 7,000 × g to pellet mitochondria. The supernatant was mixed with OptiPrep to achieve a 30% iodixanol concentration. This mixture was placed in ultracentrifuge tubes and overlaid with equal volumes of 20% and 10% iodixanol layers, then centrifuged at 100,000 × g for 2 hours. Ten fractions were collected from the top (fraction 1) to the bottom (fraction 10). Western blot analysis identified Calnexin-positive fractions, with fraction 5 showing positivity and selected for lipidomic analysis.

### Western Blot

Cells were lysed using Pierce RIPA buffer (89900, ThermoFisher) with a protease inhibitor cocktail (11697498001, cOmplete, Roche). The protein lysates were denatured by heating at 70 °C for 10 minutes in NuPAGE LDS sample buffer (NP0007, ThermoFisher) with 10 mM DTT. Proteins were separated on 4– 12% NuPAGE Bis-Tris gels (NP0336BOX, ThermoFisher) and transferred onto PVDF membranes via a semi-dry transfer at 150 mA. The membranes were blocked for 1 hour with 5% milk in PBS-T (PBS with 0.1% Tween-20), then incubated overnight at 4 °C with primary antibodies diluted in 1% milk in PBS-T. The primary antibodies used included: anti-SOAT1 (1:1000, ab39327, Abcam), anti-SREBP2 (1:1000, ab30682, Abcam), and anti-GAPDH (1:2000, AM4300, ThermoFisher). After three PBS-T washes, the membranes were incubated for 1 hour at room temperature with HRP-conjugated secondary antibodies diluted 1:10,000 in 1% milk in PBS-T (goat anti-mouse IgG, 1706516; goat anti-rabbit IgG, 1706515; Bio-Rad). Before detection, membranes underwent four additional PBS-T washes. Chemiluminescent signals were generated using the SuperSignal West Dura Extended Duration Substrate kit (34075, ThermoFisher) and visualized with an ImageQuant LAS 4000 mini system.

### Statistical analysis

Normality and homogeneity of variances were tested with the Shapiro–Wilk and Levene’s tests, respectively. For data that were normally distributed with equal variances, comparisons used a two-tailed unpaired Student’s t-test or a one-way ANOVA. If data were normally distributed but variances were unequal, Welch’s t-test was employed.

For non-parametric data or when the normality assumption was not met, group differences were evaluated using the Mann–Whitney or Kruskal–Wallis test. When multiple non-parametric groups were compared, post hoc pairwise Mann–Whitney tests with Bonferroni correction were performed.

Statistical analyses were performed on individual values (cells or GERVs) in Python using the SciPy (stats) library. Unless p-values are explicitly indicated in the figures, significance is indicated as follows: **** for p < 0.0001, *** for p < 0.001, ** for p < 0.01, * for p < 0.05, and ns for p > 0.05.

### Data representation

Graphs were generated using the Seaborn and Matplotlib libraries in Python (3.11). In most cases, SuperPlots were used to visualize biological variability and replicate consistency, with individual cells or GERV measurements shown as small points and the corresponding mean for each biological replicate shown as larger points. Data are presented as the mean of independent biological replicates ± standard deviation.

## Supplementary text

### Back to the envelope calculation of encounter times for SCAP-Cholesterol and SCAP-INSIG

The model focuses on identifying which step in the SCAP/INSIG/SREBP2 pathway is the slowest and how co-confinement within CE microdomains affects it. It calculates average first-passage times for two limited-diffusion steps, SCAP locating cholesterol and SCAP finding INSIG, using two-dimensional Smoluchowski encounter rate theory. The domain enrichment analysis uses the measured enrichment factor as input and examines how co-confinement within CE domains influences the productive encounter timescale.

All parameters are set to their most unfavorable (worst-case) values, those most challenging to the hypothesis that diffusion is not rate-limiting. Specifically, the ER membrane area is maximized, SCAP and INSIG copy numbers are halved relative to typical estimates, and diffusion coefficients are reduced by a factor of four to six compared to published measurements for similar proteins. Thus, the conclusions provide a conservative lower bound on the rate at which these encounters could occur.

### Reaction–diffusion framework for SCAP encounter kinetics in the ER membrane

The ER is the most extensive membrane system in hepatocytes, with an estimated surface area 10 to 20 times that of the plasma membrane. We use an upper bound of 40.000 µm² for the total bilayer area ^36^(see also Bionumbers and morphometric studies on mammalian cells). This is the worst-case value: a larger area distributes molecules more sparsely, increasing encounter times and making it harder to satisfy the diffusion-not-rate-limiting argument.

### Molecular densities

Molecular copy numbers were converted to area densities, assuming a homogeneous distribution across the ER membrane, using

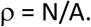

The ER contains approximately 5 mol% cholesterol relative to lipids, approximately 10^8^ molecules. Cholesterol is a small symmetrical molecule that flip-flops; it therefore equilibrates across the full bilayer thickness and is accessible to the SCAP sterol-sensing domain, which spans the transmembrane region, from both leaflets.

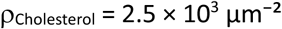

We estimate that SCAP and INSIG are present at approximately 10.000 copies per cell ^37,38^, under the worst-case assumption.

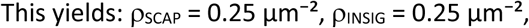

### Diffusion model

Lateral diffusion in the ER membrane was modeled as two-dimensional Brownian motion. Species-specific diffusion coefficients were assigned values lower than those obtained from experimental measurements (approximately 10 times lower for the proteins); we are making diffusion as slow as possible to yield the most conservative encounter time. D_cholesterol_ = 2µm^2^s^-1^ and D_SCAP or INSIG_ = 0.05 µm^2^s^-1^ as a conservative lower bound (we measured 0.1µm^2^s^-1^).

For encounter calculations, what matters is the relative diffusion coefficient: the effective diffusion coefficient governing the mutual approach of two independently diffusing molecules. Because both molecules move randomly and independently, their relative diffusion coefficient is simply the sum of their individual coefficients:

Pairwise effective diffusion coefficients were approximated as

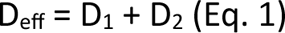

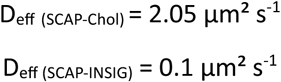

### Encounter Radii

The encounter *S* a is the center-to-center distance at which two molecules are considered to have encountered; the distance at which the relevant molecular interaction becomes accessible. It is defined as the sum of the effective radii of the two partners at the interface of interaction.

For SCAP and cholesterol: cholesterol has an effective molecular radius of approximately 0.4 nm. The SCAP sterol-sensing domain, which forms the cholesterol binding site, has an effective radius of approximately 5.0 nm based on the cryo-EM structure of SCAP.

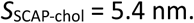

For SCAP and INSIG: both are large multi-pass membrane proteins with effective transmembrane radii of approximately 5 nm each. The encounter radius reflects the distance at which the SCAP MELADL motif is within reach of the INSIG binding interface.

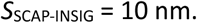

For two species diffusing independently on a 2D membrane surface, the encounter rate constant between a searcher (SCAP) and a population of targets at surface density ρ with encounter radius sigma is:

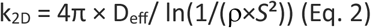

The numerator, 4π × D_eff,_ reflects how quickly the two molecules approach each other. The denominator is a logarithmic term that captures the 2D search geometry: it depends on how densely the target molecules are distributed on the membrane. A larger logarithm means a lower rate constant and a longer encounter time.

The logarithmic denominator is evaluated numerically. For the denominator not to be zero or negative, the argument of the logarithm must be greater than one, which requires ρ×*S*² to be less than one. This condition is satisfied in both cases here.

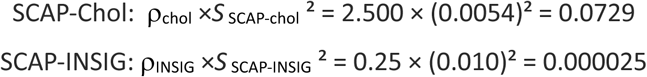

yielding

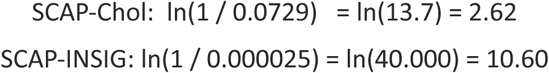

The SCAP-INSIG logarithm is larger than the SCAP-Chol logarithm because INSIG is far rarer on the membrane surface than cholesterol. A larger logarithm in the denominator reduces the rate constant and increases the encounter time, correctly reflecting that SCAP has greater difficulty finding INSIG than cholesterol does.

### Encounter rate constants

Substituting into Eq. 2 implies

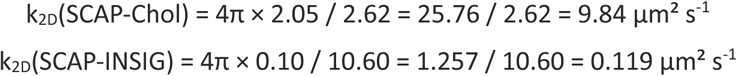

The mean first-passage time for a single SCAP molecule to encounter its nearest target is:

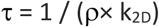

Substituting the values from Sections 2 and 5:

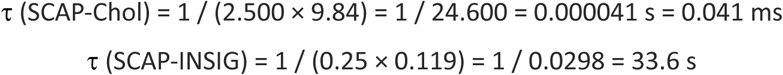

### These are the two key model outputs. SCAP finds cholesterol in 0.041 ms under worst-case conditions. SCAP finds INSIG in 33.6 s under worst-case scenario. Both are substantially faster than the rate-limiting step of productive complex formation

These values highlight that the bottleneck is the productive binding probability per encounter: the joint probability that SCAP exposes its MELADL motif, INSIG is simultaneously unoccupied and binding-competent, and both proteins meet with compatible transmembrane helix geometry.

